# Drug and siRNA screens identify ROCK2 as a therapeutic target for ciliopathies

**DOI:** 10.1101/2020.11.26.393801

**Authors:** Alice V. R. Lake, Claire E. L. Smith, Subaashini Natarajan, Basudha Basu, Sunayna K. Best, Thomas Stevenson, Rachel Trowbridge, Sushma N. Grellscheid, Jacquelyn Bond, Richard Foster, Colin A. Johnson

**Author notes:** indicates joint first authorship, these authors contributed equally to this work (Alice V. R. Lake, Claire E. L. Smith).

## Abstract

Primary cilia are microtubule-based organelles that act as cellular antennae to mediate vertebrate development and growth factor signalling. Defects in primary cilia result in a group of inherited developmental conditions known as ciliopathies. Ciliopathies often present with cystic kidney disease, a major cause of early renal failure that requires renal replacement therapies. Currently, only one drug, Tolvaptan, is licensed to slow the decline of renal function for the ciliopathy polycystic kidney disease. Novel therapeutic interventions for these conditions remain a pressing clinical need.

We screened clinical development compounds for positive effects on cilia formation and function and identified fasudil hydrochloride as the top hit. Fasudil is a generic, off-patent drug that is a potent but broadly selective Rho-associated coiled-coil-containing protein kinase (ROCK) inhibitor. In a parallel whole genome siRNA-based reverse genetics phenotypic screen of positive modulators of cilia formation, we identified ROCK2 as the target molecule. We demonstrate that ROCK2 is a key mediator of cilium formation and function through effects on actin cytoskeleton remodelling. Our results indicate that specific ROCK2 inhibitors such as belumosudil (KD-025) could be repurposed for pharmacological intervention in cystic kidney disease. We propose that ROCK2 inhibition represents a novel, disease-modifying therapeutic approach for heterogeneous ciliopathies.

## Introduction

Primary cilia are microtubule-based organelles extending from the apical surface of most mammalian cells. They act as universal cellular “antennae” in vertebrates that receive and integrate mechanical and chemical signals from the extracellular environment, serving diverse roles in chemo-, mechano- and photo-sensation (1). Cilia mediate diverse vertebrate developmental and growth factor signalling pathways, including those activated by Sonic Hedgehog (Shh), PDGF and Wnt ligands (2–4). Defects in primary cilia are associated with heterogeneous but comparatively common Mendelian inherited conditions known as the ciliopathies (5), with an overall estimated prevalence of 1 in 2000 (6). Affected children and adults commonly present with cystic kidney disease and have other clinical features such as retinal degeneration. Cystic kidney disease, associated with ciliopathies such as autosomal dominant polycystic kidney disease (ADPKD), Joubert syndrome and nephronophthisis, is a major cause of renal failure within the first two decades of life. The incidence of these ciliopathies range between 200 to 300 live births per annum in the UK (6), and we estimate that about 4000 patients in the UK require renal replacement therapy (dialysis and transplantation) that can be attributed to cystic kidney disease leading to kidney failure.

Tolvaptan, a selective vasopressin V2 receptor antagonist, slows the decline in renal function for ADPKD (7, 8). However, common (11 to 13%) side effects of Tolvaptan include polyuria, increased thirst, pollakiuria and xerostomia (9, 10) which mean that many patients do not tolerate treatment. Furthermore, there are no preventative treatments or new therapeutic interventions that may modify disease progression or the long-term outlook of ciliopathy patients with other types of cystic kidney disease. There are often several years between diagnosis and end-stage renal failure in the paediatric or juvenile age ranges, creating a window for therapeutic intervention. This has created an acute clinical need for the potential repurposing of existing drugs or the development of novel lead compounds that could treat cystic kidney disease. A recent cellular phenotype-based drug screen identified eupatilin, a plant flavonoid, which rescues ciliogenesis and ciliary transport defects caused by loss of the ciliopathy protein CEP290 (11). However, the advantages of a drug repurposing approach include existing efficacy and safety profiles for the drug, established in previous clinical trials for other indications. Drug repurposing therefore significantly reduces the time and investment required to make the move from laboratory to the clinic (12).

Here, we report a series of cellular phenotype-based compound library screens of clinical development compounds for effects on cilia formation and function. We identify fasudil hydrochloride, a potent but broadly selective Rho-associated coiled-coil-containing protein kinase (ROCK) inhibitor as the top drug hit. In a parallel whole genome siRNA-based reverse genetics phenotypic screen of positive modulators of cilia formation, we identify ROCK2 as the target molecule, and demonstrate that ROCK2 is a key mediator of cilium formation and function through downstream effects on actin cytoskeleton remodelling. Specific ROCK2 ablation or inhibition has a broad effect in rescuing ciliary function over several ciliopathy disease classes. We propose that ROCK2 inhibition represents a novel, disease-modifying therapeutic approach for heterogeneous ciliopathies. These results indicate that fasudil hydrochloride, or another more specific ROCK2 inhibitor such as belumosudil (KD025), could be repurposed for pharmacological intervention in cystic kidney disease.

## Results

### Drug screen

We selected the Tocriscreen Total library of 1120 biologically active clinical development compounds to screen for potential drugs that restore cilia after small interfering RNA (siRNA) knockdown. We carried out the screen using the mouse inner medullary collecting duct (mIMCD3) cell-line, which display primary cilia that are easy to detect, using automated imaging platforms. We reverse transfected siRNA targetting *Rpgrip1l* transcripts (si*Rpgrip1l*, designed against RefSeq NM_173431) encoding the essential ciliary protein RPGRIP1L, or a non-targetting (“scrambled”) siRNA negative control (siScr), followed by addition of chemicals (to final concentration 10 μM) after 24 hrs. We determined cell number and cilia incidence 72 hrs after transfection by imaging at three focal planes to detect the nucleus, cytoplasm and cilium (Figure 1, A, B and C). Robust *z* scores (13) were calculated to test for statistical significance in comparisons of cell number and cilia incidence (*z*_cell_ and z_cilia_). The high reproducibility of these knockdown assays was confirmed by Pearson’s correlation coefficient (R^2^) values of 0.863 and 0.781 for si*Rpgrip1l* and siScr knockdowns, respectively, for z_cilia_ between the two replicates of the screen. The strictly standardised mean difference (SSMD) for the screen, which quantifies the difference between negative and positive siRNA controls (14), had a mean value of 2.702 indicating that the siRNA effect size was “strong” (3>SSMD≥2) as defined previously (15). A transfection control (si*Plk1*) resulted in a 78.4% mean reduction in cell number compared to negative controls (mean *z*_cell_ −57.36) indicating that siRNA reverse transfection was highly efficient (Figure 1, D and E). Transfection of positive controls si*Ift88* and si*Rpgrip1l* resulted in significant decreases in cilia incidence (percentage of cells with a single cilium reduced by 9.8% and 17.1%, z_cilia_ −7.67 and −12.96, respectively) compared to negative controls.

**Figure 1:**
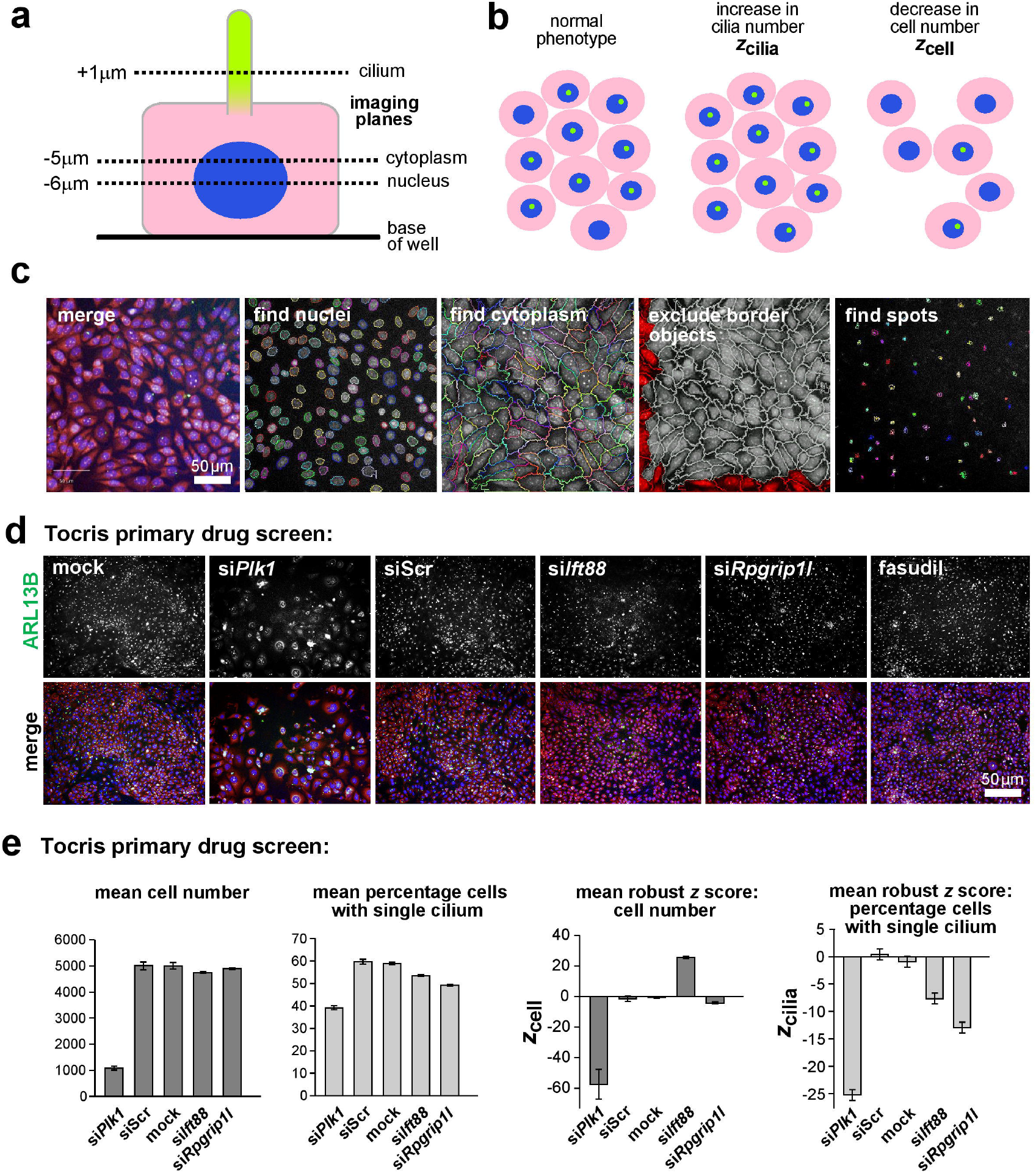
High content imaging protocol for high throughput ciliogenesis screening. **(a)** Schematic of a polarised mIMCD3 cell showing the focal planes used to image nuclei (blue), cytoplasm (pink) and ciliary axonemes (green). **(b)** Robust *z* scores were calculated to identify significant changes in cilia incidence (*z*_cilia_) or in cell number (*z*_cell_) **(c)** mIMCD3 cells imaged using an Operetta high-content imaging system. Merge images show staining for cilia marker ARL13B (green), nuclei (DAPI; blue) and cellular RNA (TOTO-3; pink). Representative images are also shown from Harmony/Columbus software of cell segregation and cilia recognition (“find spots”) protocols. Scale bar = 50 μm. **(d)** Representative immunofluorescence high content images from the Tocris primary drug screen showing decrease of cilia incidence following reverse transfection with positive control siRNA pools against *Plk1, Ift88* and *Rpgrip1l* compared to the non-targeting scrambled siRNA (siScr). Following knockdown with si*Rpgrip1l*, cilia incidence was rescued with 10 μM fasudil hydrochloride. Scale bar = 50 μm. **(e)** Left: bar graphs quantitate the effect on cell number and ciliogenesis (mean % cells with a single cilium) for positive controls (*Plk1, Ift88* and *Rpgrip1l* siRNAs) and negative controls (siScr and mock transfections). Right: bar graphs showing mean robust *z* scores for cell number (*z*_cell_) and mean robust *z* score for cilia incidence (*z*_cilia_) for positive and negative siRNA controls. Error bars indicate s.e.m. for n=8 experimental replicates.

For each screening batch (corresponding to four 96-well plates within the two replicates), we normalized robust *z* scores across the screen in order to set cut-off values so that we could select hits within each batch (Figure 2, A, B and C). Hits were selected if *z*_cilia_ ≥ *z*_ciliacutoff_ (where *z*_ciliacutoff_ = normalized *z*_cilia_ of batch controls + 2). Chemicals were excluded from further analysis if *z*_cell_ ≤ *z*_cellcutoff_ (where *z*_cellcutoff_ = normalized *z*_cell_ of batch controls - 2) for >2 of the 4 plates in each batch, in order to remove those that had a significant cytotoxic effect. Following the normalization of cut-off values, we selected 73 chemicals for secondary screening, although only 71 were commercially available (Supplemental Data 1). Secondary screening in mIMCD3 cells identified 25 hit compounds (Figure 2, D). We then did a tertiary screen in a cell-line of a different species (the human ciliated cell-line hTERT RPE-1), using siRNAs of a different chemistry against human *RPGRIP1L* (RefSeq NM_015272) and an additional independent ciliary target (human *IFT88* RefSeq NM_175605, encoding the essential ciliary intraflagellar transport protein IFT88). We also used new, freshly prepared, commercial batches of each chemical. The tertiary screen validated two chemicals, fasudil hydrochloride and (RS)-4-carboxy-3-hydroxyphenylglycine (Figure 2, E). Fasudil hydrochloride (hereafter referred to as fasudil) increased cilia incidence despite knockdowns of *IFT88* and *RPGRIP1L* (*z*_cilia_ increases of +1.92 and +1.79, respectively, compared to negative controls) without any cytotoxic effect (no significant decrease in *z*_cell_ values). Negative control cells transfected with siScr and treated with fasudil also had no significant differences in either *z*_cell_ or *z*_cilia_ suggesting that fasudil was not cytotoxic and did not affect cilia incidence in normal ciliated cells. (RS)-4-carboxy-3-hydroxyphenylglycine increased cilia incidence (*z*_cilia_ increases of +0.82 and +1.04 for si*IFT88* and si*RPGRIP1L*, and +1.48 for siScr) without any significant effects on *z*_cell_. However, since the effect on cilia incidence was less significant than fasudil, (RS)-4-carboxy-3-hydroxyphenylglycine was not taken forward for dose response testing.

**Figure 2:**
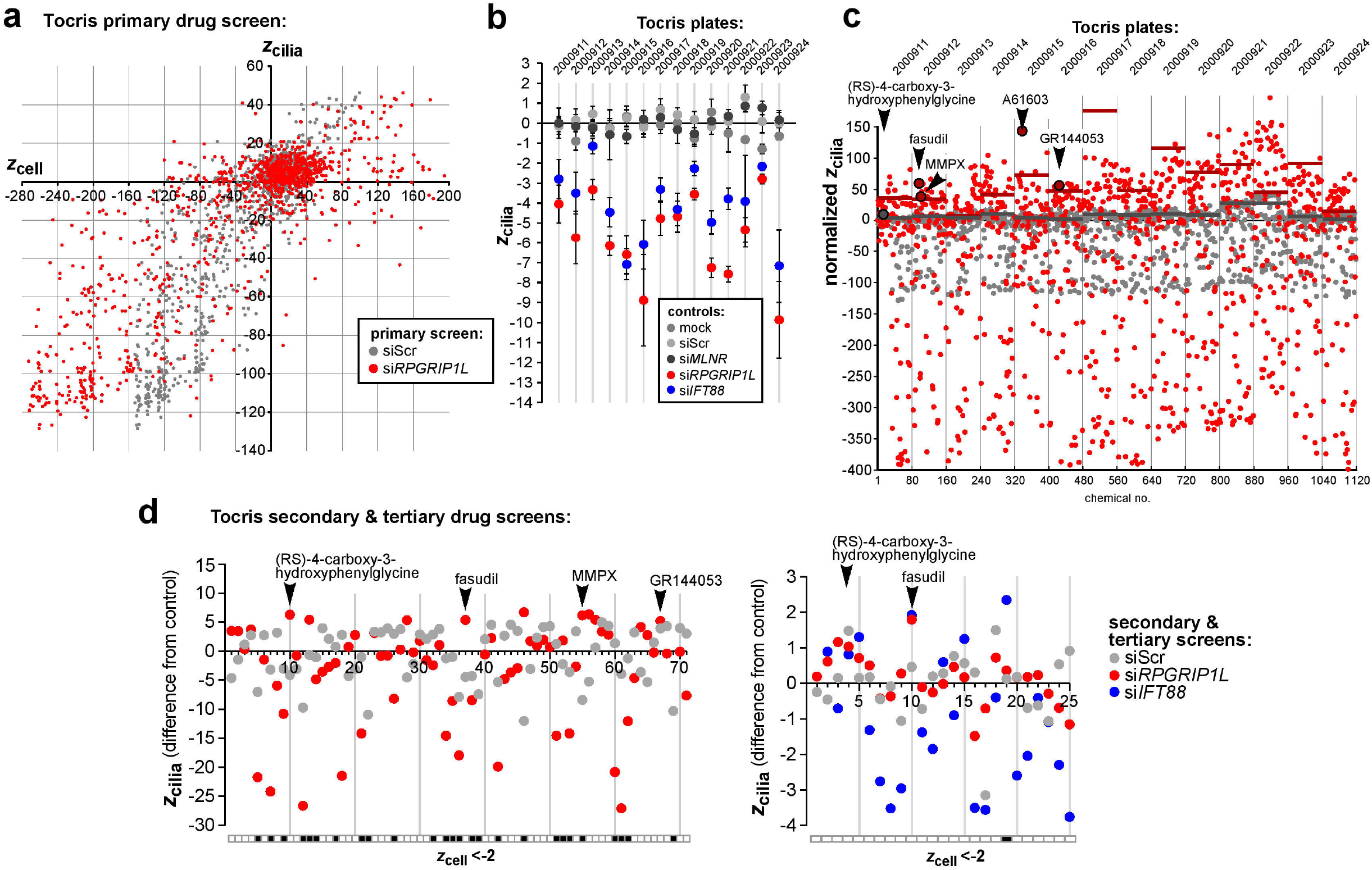
High throughput screens of cilia incidence using the Tocriscreen Total library of active clinical development compounds. **(a)** Primary drug screen summarising mean *z*_cell_ and *z*_cilia_ values (n=2) for mouse mIMCD3 cells treated with each chemical and reverse transfected with either si*Rpgrip1l* (red) or siScr (grey). **(b)** Normalised *z*_cilia_ values for cells treated with each chemical and reverse transfected with either si*Rpgrip1l* (red) or siScr (grey). Tocris plate numbers are indicated along the x-axis. Normalized *z*_cilia_ cut-off values (coloured bars) for each plate are indicated (coloured bars). Hits are defined as chemicals with normalized *z*_cilia_ greater than cut-off values. Selected hits are indicted. **(c)** Summary of negative (grey) and positive (red & blue) control *z*_cilia_ values for each Tocris plate. Error bars indicate s.e.m. for n=2 experimental replicates, each with 4-8 technical replicates, depending upon the control. All plates passed quality control criteria of *z*_cilia_ ≤ −2 for positive controls (si*Ift88* and si*Rpgrip1l*) and −2 ≤ *z*_cilia_ ≤ +2 for negative controls (mock, siScr, si*MLNR*). **(d)** Secondary screen (n=2) of 71 chemicals in mIMCD3 cells. Hits are defined as chemicals with *z*_cilia_ difference from relevant control of ≥ +2. Hits were excluded if *z*_cell_ ≤ −2 (black filled squares in the grid). **(e)** Tertiary screen (n=2) of 25 chemicals in human hTERT RPE-1 cells. Hits are defined as chemicals with *z*_cilia_ difference from relevant control of ≥ 1.5. Hits were excluded if *z*_cell_ ≤ −2 (black filled squares in the grid).

### Dose response testing of fasudil

Fasudil (also known as HA-1077 and AT877) is an isoquinolone derivative based on the core structure of 5-(1,4-diazepan-1-ylsulfonyl)isoquinoline, and is a potent but broadly selective Rho-kinase (ROCK) inhibitor of both ROCK1 and ROCK2 isozymes. Fasudil is currently unlicensed by the Federal Drugs Agency (FDA) and the European Medicines Agency (EMA), but is in clinical use in Japan and China and in trials elsewhere as a treatment for cerebral vasospasm (16), pulmonary hypertension (17), to reduce cognitive decline in stroke patients (18) and for amyotrophic lateral sclerosis ALS (19).

First, we tested the effect of fasudil on cell viability, performing MTT assays in both mIMCD3 and hTERT RPE-1 cell-lines (Supplemental Figure 1, A). For mIMCD3 cells, the mean metabolic activity ranged between 81.6 to 96.9% compared to vehicle control across all time points for the range of concentrations tested. hTERT RPE-1 cells were more sensitive to the cytotoxic effects of fasudil, with mean metabolic activity that ranged from 66.9% at 100 μM to a maximum of 93.3%. Next, following siRNA knockdowns, we tested dose responses in both mIMCD3 and hTERT RPE-1 cell-lines for fasudil concentrations ranging from 1 μM to 100 μM (Supplemental Figure 1, B). We confirmed significant rescue of cilia incidence following either si*Ift88* or si*Rpgrip1l* knockdowns even with 1 μM fasudil, the lowest concentration tested. Significant rescue was maintained up to 30 μM, but with significant cytotoxic effects (*z*_cell_ < −2) at concentrations ? 30 μM. Dose response assays in hTERT RPE-1 cells showed significant rescue at 10 μM and higher fasudil concentrations for si*RPGRIP1L*-treated but not for si*IFT88*-treated cells, although cytotoxicity was apparent at lower concentrations than for mIMCD3 cells.

### Fasudil can restore cilia in CRISPR-Cas9 edited cell-lines but other derivatives are cytotoxic

We then investigated the effects of fasudil in alternative cellular models of ciliopathies. We used CRISPR-Cas9 editing to create heterozygous knockout lines *IFT88^+/-^, RPGRIP1L^+/-^* and *TMEM67*^+/-^, as well as a biallelic knockout line *TMEM67*^-/-^. The null mutations were: *IFT88* (NM_175605.5):[c.371_408delinsAAGAAAAAAG,p.(P124Qfs*15)], *RPGRIP1L* (NM_015272.5):[c.15_37del, p.(D6Cfs*35)], *TMEM67* (NM_153704.6): [c.369delC, p.(E124Kfs*12)] and *TMEM67* (NM_153704.6):[c.519delT, p.(C173Wfs*20)];[c.519dupT, p.(E174*)]. We confirmed reduced expression of IFT88, RPGRIP1L and TMEM67 in each cell-line respectively by western blot or by immunofluorescence (Supplemental Figure 2). In dose response assays, cilia incidence was rescued in the *IFT88*^+/-^ and *RPGRIP1L*^+/-^ celllines at 1 μM fasudil, the lowest concentration tested, but there were significant cytotoxic effects ≥ 10 μM (Figure 3, A). In wild-type hTERT RPE-1 cells, cilia incidence was significantly rescued at 3 μM and higher concentrations but was less than that observed for the *IFT88*^+/-^ and *RPGRIP1L*^+/-^ cell lines. In contrast, there was no significant rescue of cilia incidence in either *TMEM67*^+/-^ or *TMEM67*^-/-^ cell lines for concentrations of fasudil that were not also significantly cytotoxic (>10 μM).

**Figure 3:**
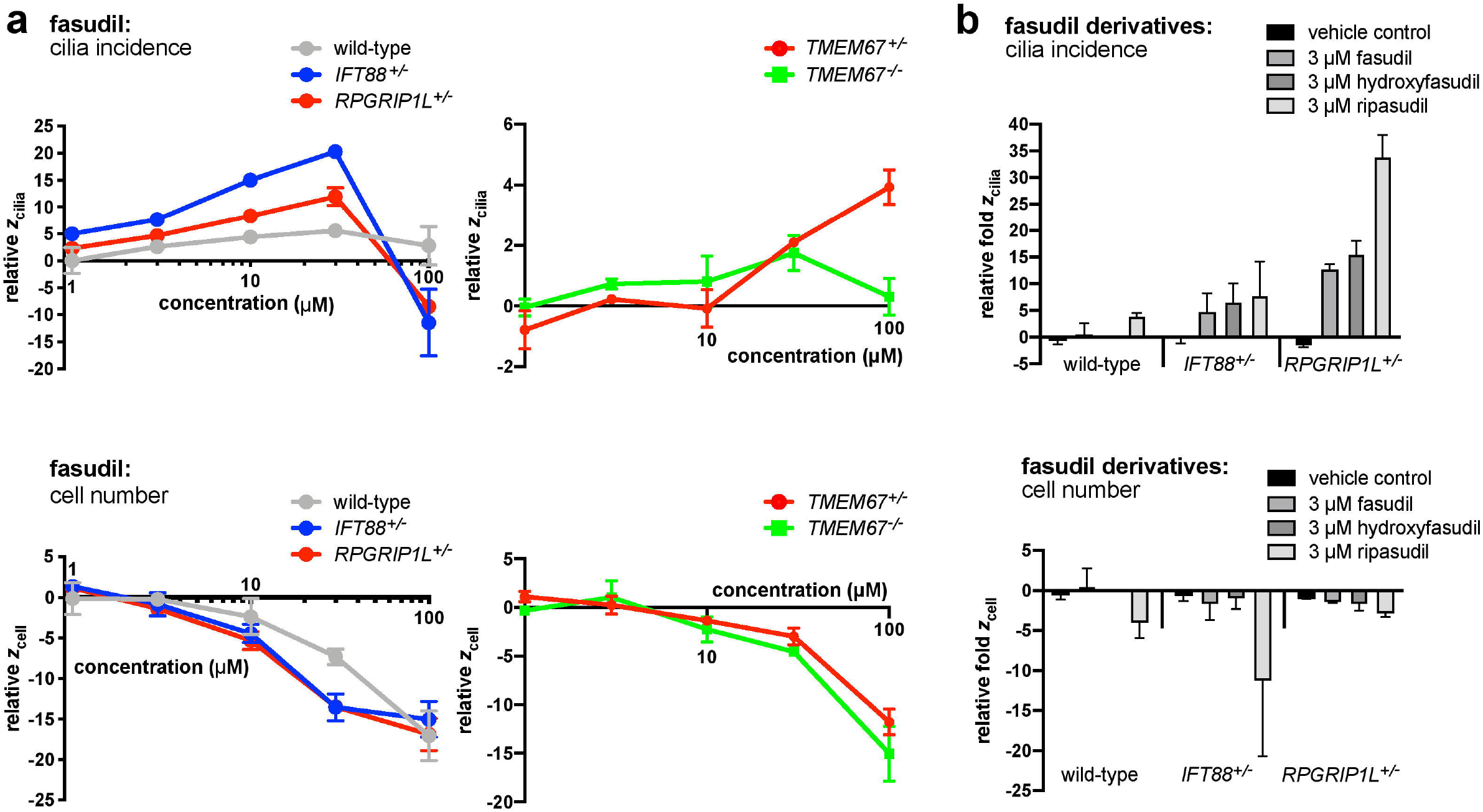
Fasudil and fasudil derivative dose response assays in wild-type and ciliopathy crispant hTERT RPE-1 cells. **(a)** Dose response assays in wild-type mother line, heterozygous *IFT88*^+/-^, *RPGRIP1L*^+/-^ and *TMEM67*^+/-^ knockout hTERT RPE-1 lines, as well as a biallelic knockout line for *TMEM67*^-/-^, following treatments with a concentration range of 1 to 100 μM fasudil. Details of cell-line genotypes are given in the main text. Graphs show the mean robust *z* scores (n=2 experimental replicates, each with n=3 technical replicates) for cilia incidence (*z*_cilia_; upper panels) and cell number (*z*_cell_); lower panels). Values are normalized to vehicle control (DMSO) treated cells. Error bars represent the range. **(b)** Dose response assays in wild-type mother line, and heterozygous *IFT88*^+/-^ and *RPGRIP1L*^+/-^ knockout hTERT RPE-1 lines for 3 μM fasudil and derivatives hydroxyfasudil and ripasudil. Bar graphs show the fold-change in *z*_cilia_ (top) *z*_cell_ (bottom) relative to vehicle control (n=2 experimental replicates, each with n=3 technical replicates). Error bars represent the range. § indicates omitted data due to technical artefact.

Fasudil is an inhibitor of both ROCK isozymes, ROCK1 and ROCK2, with a half maximal inhibition concentration (IC_50_) ranges of 0.84-2.67 and 0.69-1.90 μM, respectively (20–22) (Supplemental Table 1). However, fasudil has off-target effects on other kinases (e.g. >75% inhibition of MAPK15, RPS6KA1, PKN2 and RPS6KA at 10 μM (23), which may contribute to the cytotoxicity observed at high concentrations in our dose response testing. We investigated two other derivative compounds based on 5-(1,4-diazepan-1-ylsulfonyl)isoquinoline, hydroxyfasudil and ripasudil. Ripasudil is currently in clinical use as a topical treatment for glaucoma and ocular hypertension (24, 25). Hydroxyfasudil (also known as HA-1100) has IC_50_ values of 0.73 μM and 0.72 μM for (human) ROCK1 and ROCK2, respectively (20). Hydroxyfasudil has a more favourable kinase inhibition profile (23) that may reduce off-target effects in comparison to fasudil. In contrast, ripasudil (K115) is a more selective inhibitor of ROCK2, with IC_50_ of 0.051 μM for ROCK1 and 0.019 μM for ROCK2 (26). Both hydroxyfasudil and ripasudil significantly increased cilia incidence relative to fasudil, in both the *IFT88*^+/-^ and *RPGRIP1L*^+/-^ CRISPR-Cas9 edited cell-lines (Figure 3, B). However, ripasudil was significantly more cytotoxic at 3 μM and was therefore not considered for further analysis. We note that hydroxyfasudil was included as a clinical development compound in the primary drug screen but was excluded from secondary screening due to effects on cell number.

### Functional effects of fasudil on acto-myosin contraction and Sonic Hedgehog signalling

Since actin remodelling pathways are activated by ROCK and are known to be important in ciliogenesis and cilia maintenance (27, 28), we hypothesised that ROCK inhibition by fasudil might result in visible changes to the actin cytoskeleton. We assessed the effects of fasudil treatment on F-actin using automated high content imaging of mIMCD cells stained with phalloidin. Image structure analysis was used to quantify F-actin structures. There was no significant difference in phalloidin-stained cytoplasmic F-actin structure for cells treated with fasudil compared with vehicle control (data not shown). We also looked for possible changes in acto-myosin contractions by assessing levels of phosphorylated non-muscle myosin light chain 2 (MLC), which reflects the active state of non-muscle myosin IIA and capacity for acto-myosin contractility. Cells were stained with anti-phospho-MLC antibodies for monophosphorylated MLC IIa (Ser19) (p-MLC) and biphosphorylated MLC (Thr18/Ser19) (pp-MLC). Automated high content imaging and image structure analysis allowed quantification of fibre-like structures associated with acto-myosin contractility. Levels of pp-MLC were lower, but not significantly so, (*p*=0.055, unpaired, two-tailed Student’s t-test, n=3) in fasudil-treated cells compared with cells treated with vehicle control. Levels of MLC and p-MLC were similar in both conditions (Supplemental Figure 3).

To test if primary cilia restored by fasudil treatment are functional, we assessed the effect of this compound on Sonic Hedgehog (Shh) signalling, a key developmental signal transduction pathway that is mediated by the ciliary sub-cellular compartment. First, we measured responsiveness to Shh activation in the murine NIH3T3 “Shh-Light II” reporter cell-line. This contains the firefly luciferase reporter gene under the control of the Gli promoter. We reverse transfected the Light II cells with si*Ift88* or si*Rpgrip1l* against ciliary targets. Knockdown of Patched1 (Ptch1), the Shh receptor, was included as a positive control since this stimulates the pathway by releasing Smoothened (Smo), the downstream G protein-coupled receptor, from the ciliary membrane (29). As an additional positive control, treatment with Smoothened agonist (SAG) exacerbated Shh responses in in all knockdown conditions (**** *p*<0.0001, unpaired, two-tailed Student’s t-test, n=3), indicating that LightII cells retained correct Shh responsiveness. As expected, knockdown with si*Ift88* decreased whereas si*Ptch1* significantly increased Shh signalling (p=0.161 and *** 0.000194 respectively, unpaired, two-tailed, Student’s t-tests, n=3) (Figure 4, A). Knockdown with si*Rpgrip1l* significantly increased Shh signalling (*** p=0.000283, pairwise, two-tailed Student’s t-test, n=3), consistent with previous studies in the *Rpgrip1l*^-/-^ knockout mouse (30). We then assessed the effect of 10 μM fasudil, in both unstimulated and SAG-stimulated cells compared with treatment with vehicle control, for each knockdown condition. Treatment with 10 μM fasudil significantly and consistently increased Shh signalling in all knockdown conditions when compared to the vehicle control, both with and without SAG stimulation (si*Ift88*: * 0.0317, si*Rpgrip1l*: * 0.0118, si*Ptch1*: ** 0.00914, siScr: * 0.0441; pairwise, two-tailed Student’s t-tests, n=3). This suggests that fasudil can partially restore cilia function, as measured by ciliary-mediated Shh signalling. Second, as an independent assay to confirm this effect, we determined Smo occupancy in the ciliary membrane by using confocal microscopy. Smo accumulates in the ciliary membrane when Ptch1 is inactivated and is subsequently removed upon binding of Shh ligand to Ptch1 (31). Application of SAG resulted in Smo accumulation within cilia (Figure 4, B). Knockdown with si*Ift88* or si*Rpgrip1l* in mIMCD3 cells resulted in a significant reduction in the area of Smo localized to the cilia remaining under these knockdown conditions (**** p<0.0001, pairwise, two-tailed Student’s t-tests, n=4), indicating a decrease in Shh signalling (Figure 4, C). However, treatment with 10 μM fasudil resulted in a significant increase of Smo occupancy at cilia (** p = 0.00208, pairwise, two-tailed Student’s t-tests, n=4) confirming that fasudil can restore aspects of ciliary function in a cellular model of the ciliopathy disease state.

**Figure 4:**
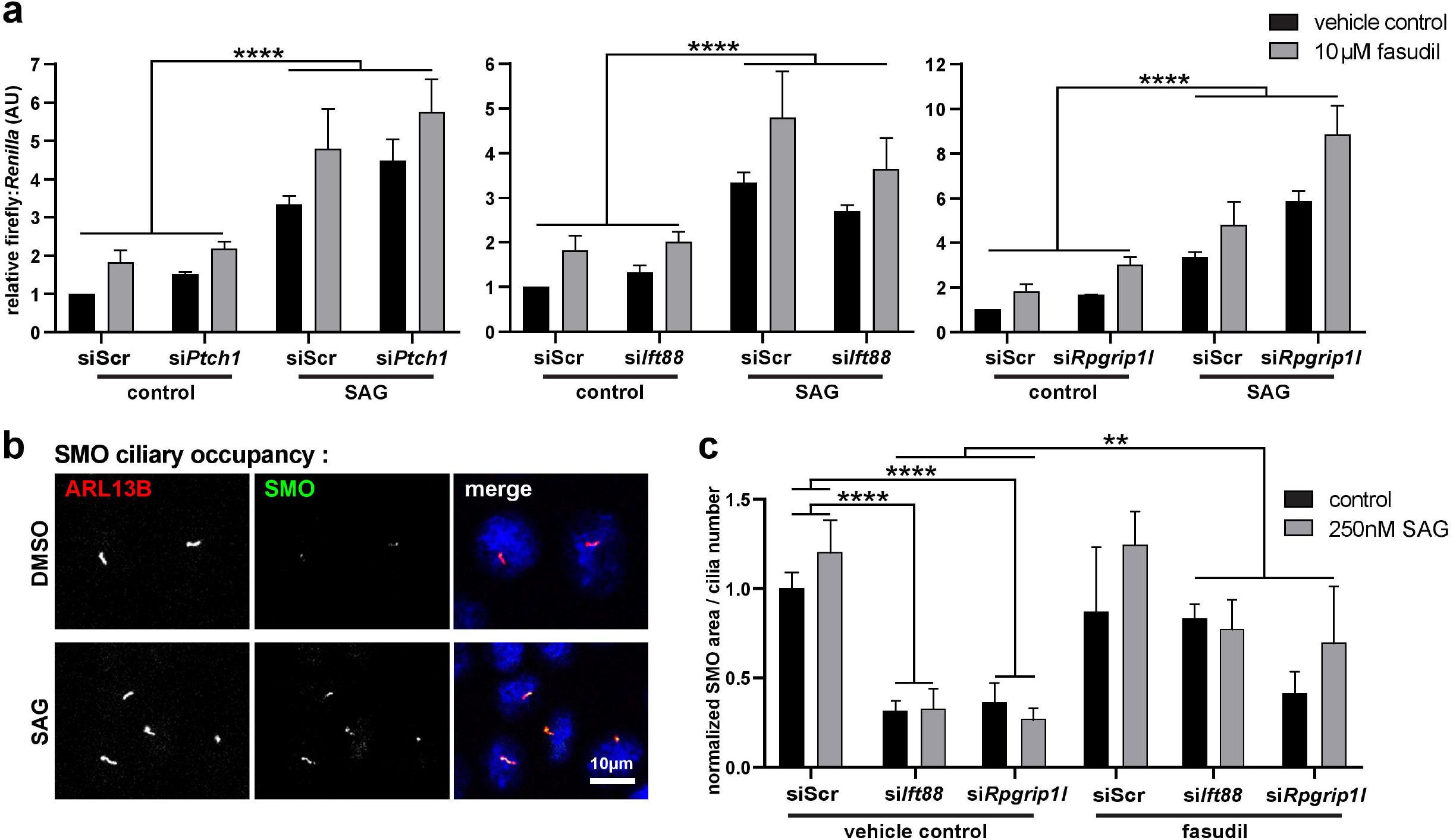
Fasudil restores ciliary-mediated Shh signalling. **(a)** Bar graphs of dual (firefly & *Renilla*) luciferase Gli reporter assays showing the effects of si*Ptch1*, si*Ift88* and si*Rpgrip1l* knockdown compared to siScr in mouse Shh-Light II NIH3T3 cells (n=3). Cells were treated with 250 nM Smoothened agonist (SAG) and/or 10 μM fasudil and compared with vehicle (DMSO) controls, with data normalized to controls transfected with siScr. SAG significantly increased the relative firefly:*Renilla* luciferase ratios in all knockdown conditions (**** p<0.0001, paired two-tailed Student’s t-test). 10 μM fasudil significantly increased the luciferase ratio in all knockdown conditions when compared to the vehicle control, both with and without SAG stimulation (si*Ift88* * *p*=0.0317, si*Rpgrip1l* * *p*=0.0118, si*Ptch1* ** *p*=0.00914). Error bars represent s.e.m. **(b)** Representative confocal immunofluorescence microscopy images of mIMCD3 cells treated with SAG or vehicle control. Cilia were stained for ARL13B (red) and Smoothened (SMO; green). SAG treatment resulted in qualitatively more SMO within the ARL13B-marked cilia. Scale bar=10 μm. **(c)** Bar graph shows the normalized area of SMO localization per cilium for mIMCD3 cells knocked down with siScr, si*Ift88* or si*Rpgrip1l*. Knockdown with si*Ift88* or si*Rpgrip1l* significantly reduced the area of SMO localized to each cilium when treated with vehicle control (**** p<0.0001). Treatment with 10 μM fasudil resulted in a significant increase in SMO area per cilium (** *p*=0.00208). Statistical significance was calculated with paired twotailed Student’s t-tests (n=2 experimental replicates, each with n=2 technical replicates, minimum of 19 cilia analysed per replicate. Error bars represent s.e.m.

### Secondary siRNA screening identified ROCK2 as a negative regulator of cilia incidence

We aimed to determine key molecular drug targets that could increase cilia incidence, which could also represent the main molecular effector of fasudil. We previously published a genome-wide siRNA reverse genetics screen in mIMCD3 cells (32) and have re-analysed the raw image data to identify hits that increased cilia incidence. The screen data was filtered for hits that: i) significantly increased cilia incidence *z*_cilia_ > +2; ii) did not have any significant effect on cell number +2> *z*_cell_ > −2; iii) used an siRNA pool that targeted all annotated transcripts of the gene; and iv) had a human orthologue so that follow-up investigations could be carried out (Figure 5, A). We identified 83 hits from the re-analysed primary screen that were taken forward for secondary screening using the same screening methods. Eight of 83 hits significantly increased cilia incidence across replicates of the secondary screen (Figure 5, B, Supplemental Data 2). Positive (si*Plk1*, si*Rpgrip1l* and si*Ift88*) and negative controls (siScr and mock transfections) verified that our assay was robust and could identify targets that caused changes in cell numbers and cilia incidence (Figure 5, C). Comparison of the two replicates of the screen showed that the mean R^2^ values between runs was 0.6366 for cell number and 0.7156 for cilia incidence, indicating high reproducibility of the assay (data not shown). The top hit from the secondary screen was *Rock2*, with an average *z*_cilia_ score of +3.796 and *z*_cell_ of −0.300. Images from the secondary screen also qualitatively show that cilia incidence is increased compared to the siScr control (Figure 5, D). Tertiary screening validated *ROCK2* as a hit following knockdown in the human hTERT RPE-1 cell-line using siRNAs of a different chemistry (Supplemental Data 2), and efficient knockdowns by both human and mouse siRNAs were confirmed by western blots (Figure 6, A, B). *ROCK2* knock-down in hTERT RPE-1 cells increased cilia incidence by >10% and significantly increased average cilia length (by 34.7%) (Figure 6, C). Over-expression of GFP-tagged m*Rock2* significantly decreased both cilia incidence and cilia length compared to controls, consistent with a dominant negative effect (Figure 6, D). These results suggest that ROCK2 has a conserved role in mammals during ciliogenesis.

**Figure 5:**
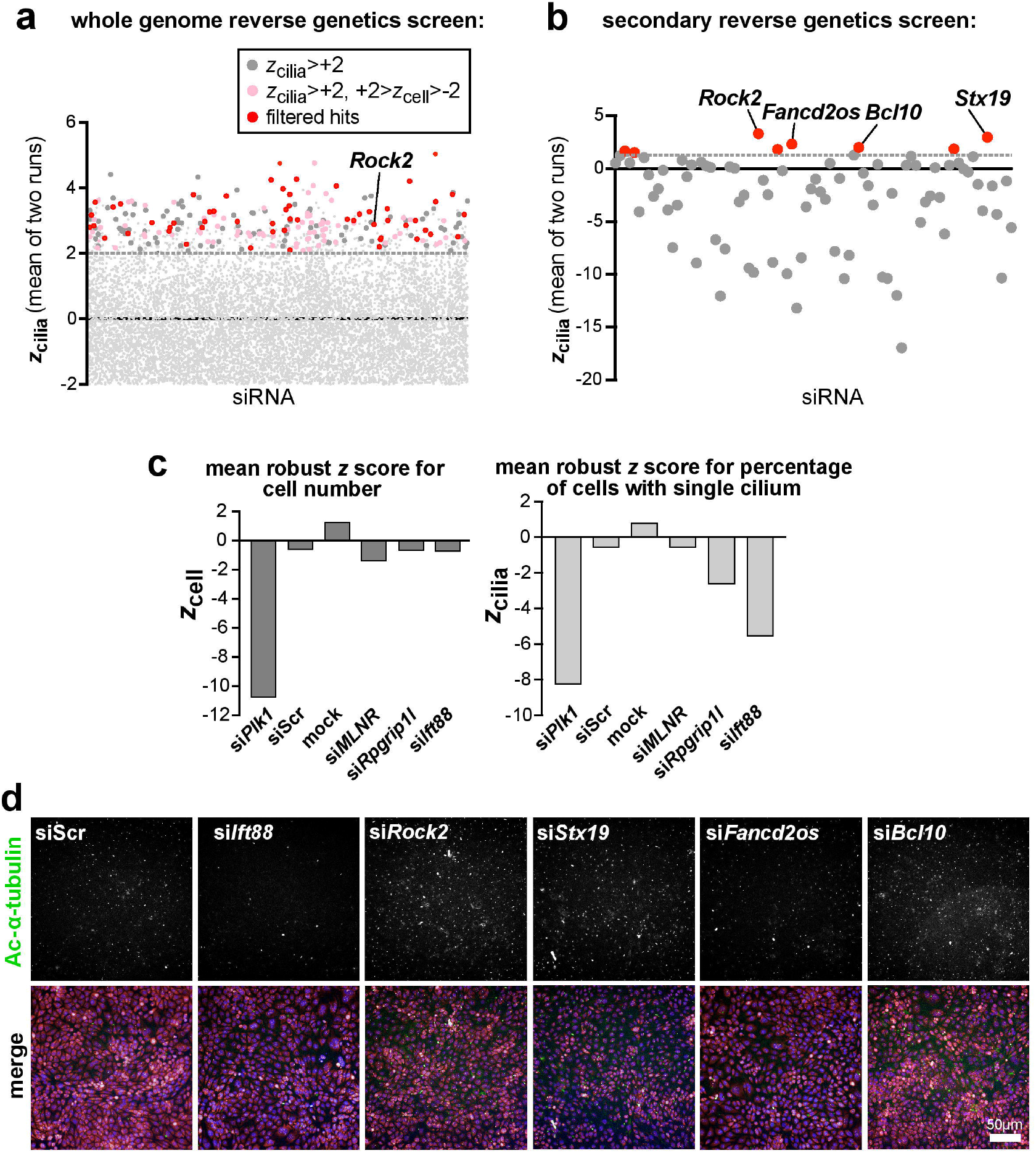
Whole genome siRNA reverse genetics screen of cilia incidence. **(a)** Whole genome siRNA primary reverse genetics screen summarising mean *z*_cilia_ values (n=2) for mouse mIMCD3 cells; 8907 data points are outside of the y-axis limit of −2.0. Dark grey points indicate hits *z*_cilia_ ≥ +2 (cut-off indicated by grey dashed line), and pink points indicate hits with no significant effect on cell number (−2 ≤ *z*_cell_ ≤ +2). Red points indicate 83 hits taken forward for secondary screening that have human orthologues and siRNAs targeting all transcripts of the gene. **(b)** Secondary screen (n=2) of 83 primary screen hits in mIMCD3 cells, validating 8/83 hits (red points) with mean *z*_cilia_ ≥ +2. The top four hits are indicated with *z*_cilia_ values as follows: *Rock2* +3.80, *Stx19* +3.46, *Fancd2os* +2.83, *Bcl10* +2.49. **(c)** Bar graphs showing mean *z*_cell_ (left) and *z*_cilia_ (right) for positive controls (*Plk1, Ift88* and *Rpgrip1l* siRNAs) and negative control siRNAs (siScr and mock transfections). Error bars indicate s.e.m. for n=8 experimental replicates. **(d)** Representative immunofluorescence high content images from the primary screen for top hits (*siRock2, siStx19, siFancd2os* and *siBcl10* knockdowns) showing increase in cilia incidence compared to scrambled siRNA (siScr) negative control; si*Ift88* is included as a positive control for cilia loss. Merge images show staining for cilia marker acetylated α-tubulin (green), nuclei (DAPI; blue) and cellular RNA (TOTO-3; pink). Scale bar = 50 μm.

**Figure 6:**
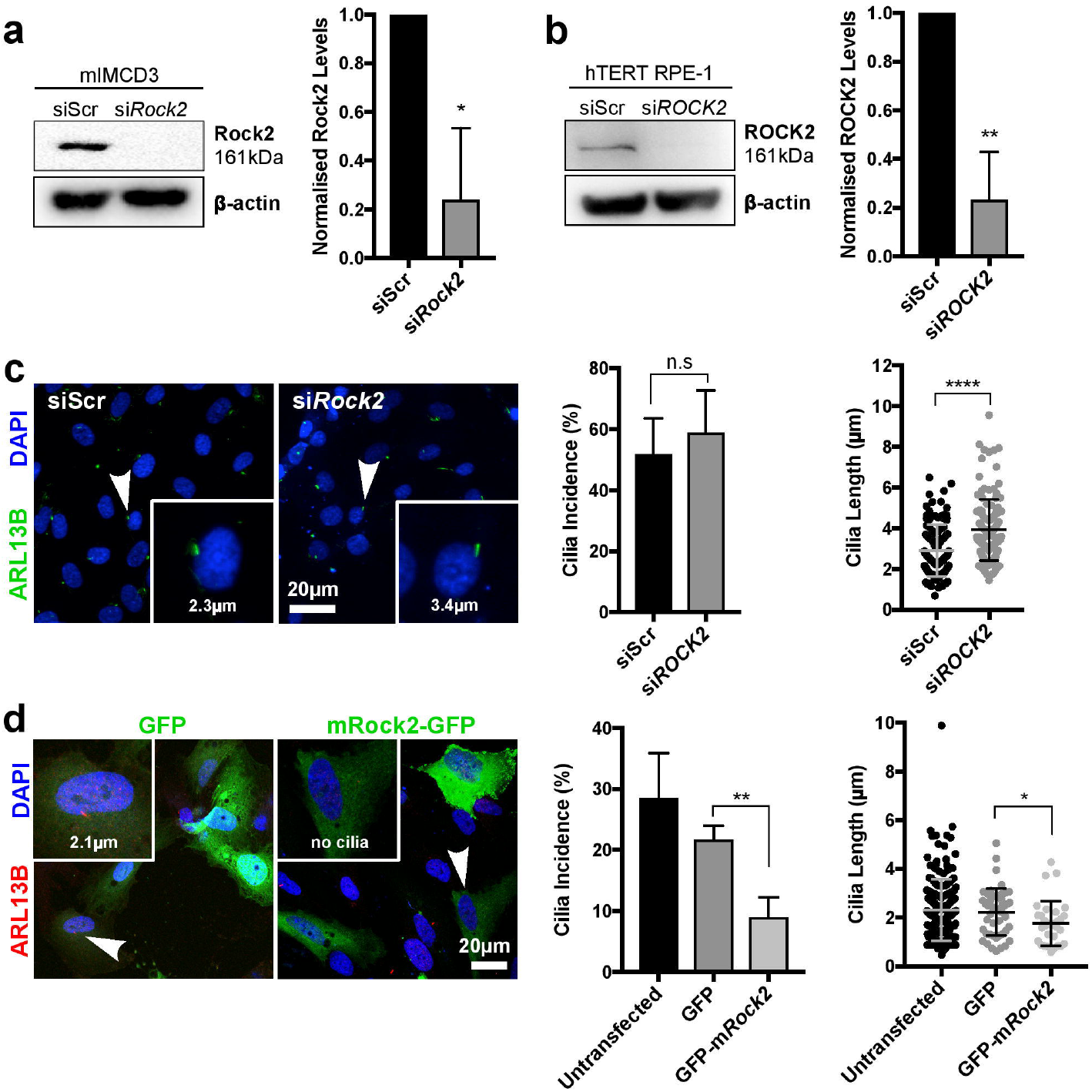
ROCK2 is a negative modulator of ciliogenesis. Validation of siRNA knockdowns for murine **(a)** and human **(b)** ROCK2. Western blots indicate loss of ROCK2 following knockdowns in mIMCD3 and hTERT RPE-1 cells compared to scrambled siRNA (siScr) negative control. Bar graphs quantify western blot densitometry measures (n=3) for each cell line, normalised against β-actin levels. Statistical significance of pairwise comparisons are indicated (* *p*<0.05, ** *p*<0.01; unpaired two-tailed Student’s t-test). **(c)** Left: immunofluorescence confocal microscopy of hTERT RPE-1 cells, immunostained for cilia marker ARL13B (green), after knockdown with *siROCK2* and showing increase in cilia length. Arrowheads indicate cilia displayed in magnified insets, shown with their measured lengths. Right: bar graphs showing mean cilium length for siScr treatment was 2.91 μm, compared to si*ROCK2* knockdown 3.92 μm (n=3, minimum of 40 cilia measured per replicate). Statistical significance of pairwise comparison is indicated (**** *p*<0.0001 unpaired two-tailed Student’s t-test). There was no significant difference in cilia incidence (n=3, *p*=0.5310). Scale bar = 20 μm. **(d)** Left: confocal microscopy of hTERT RPE-1 cells transiently transfected with either untagged GFP or GFP-m*Rock2* constructs (green) and stained for ARL13B (red). Arrowheads indicate cells displayed in magnified insets. Right: bar graphs showing that cells transfected with GFP-m*Rock2* had a significantly lower cilia incidence (p=0.0049) and shorter cilia (p=0.012) compared to cells expressing untagged GFP (n=3). Statistical significance of pairwise comparisons calculated with an unpaired twotailed Student’s t-test (cilia incidence) and Mann-Whitney U-test (cilia length) as indicated (* *p*<0.05, ** *p*<0.01). Scale bar = 20 μm.

### *ROCK1 cannot compensate for loss of ROCK2* function during ciliogenesis

Rock1 (the isozyme of Rock2, with 93% kinase domain homology) (33), did not pass the initial stringent filtering of primary whole genome siRNA screen data (mean *z*_cilia_ = +0.864), although siRNA knockdown had a marginal positive effect on cilia incidence that is consistent with the results in a previous siRNA screen of “druggable targets” (34). Since fasudil is a broadly selective inhibitor of both ROCK isozymes, it was important to determine which isozyme had the predominant effect on ciliogenesis since they both have distinct and separate functional roles in other cellular mechanisms (35–38). We hypothesised that ROCK1 has a redundant role in ciliogenesis because residual ROCK1 activity had not been able to compensate for the loss of ROCK2 in our previous experiments. Efficient knockdowns by both mouse and human siRNAs against *ROCK1* were confirmed by western blots (Supplemental Figure 4, A). We then determined that knock-downs of *ROCK1* in either mIMCD3 or hTERT RPE-1 cells had no significant effects on cilia incidence (Supplemental Figure 4, B, C), but did increase cilia length in mIMCD3 cells. This suggests that the function of ROCK2 during ciliogenesis is non-redundant and is not fully compensated by ROCK1.

To further validate this finding, we used the specific ROCK2 inhibitor belumosudil (also known as KD025 and SLX-2119) which has IC_50_ values of 24 μM and 0.105 μM for ROCK1 and ROCK2, respectively (Supplemental Table 1). Belumosudil treatment over a 2hr period caused moderate but significant increases in cilia incidence and cilia length (Supplemental Figure 5, A, B). However, more significant effects were observed in both mIMCD3 and hTERT RPE-1 (Figure 7, A, Supplemental Figure 5, B) cells following treatment with belumosudil for 48hrs. This suggested that the inhibition of ROCK2 was the predominant effect leading to increased ciliogenesis and rescue of cilia function.

**Figure 7:**
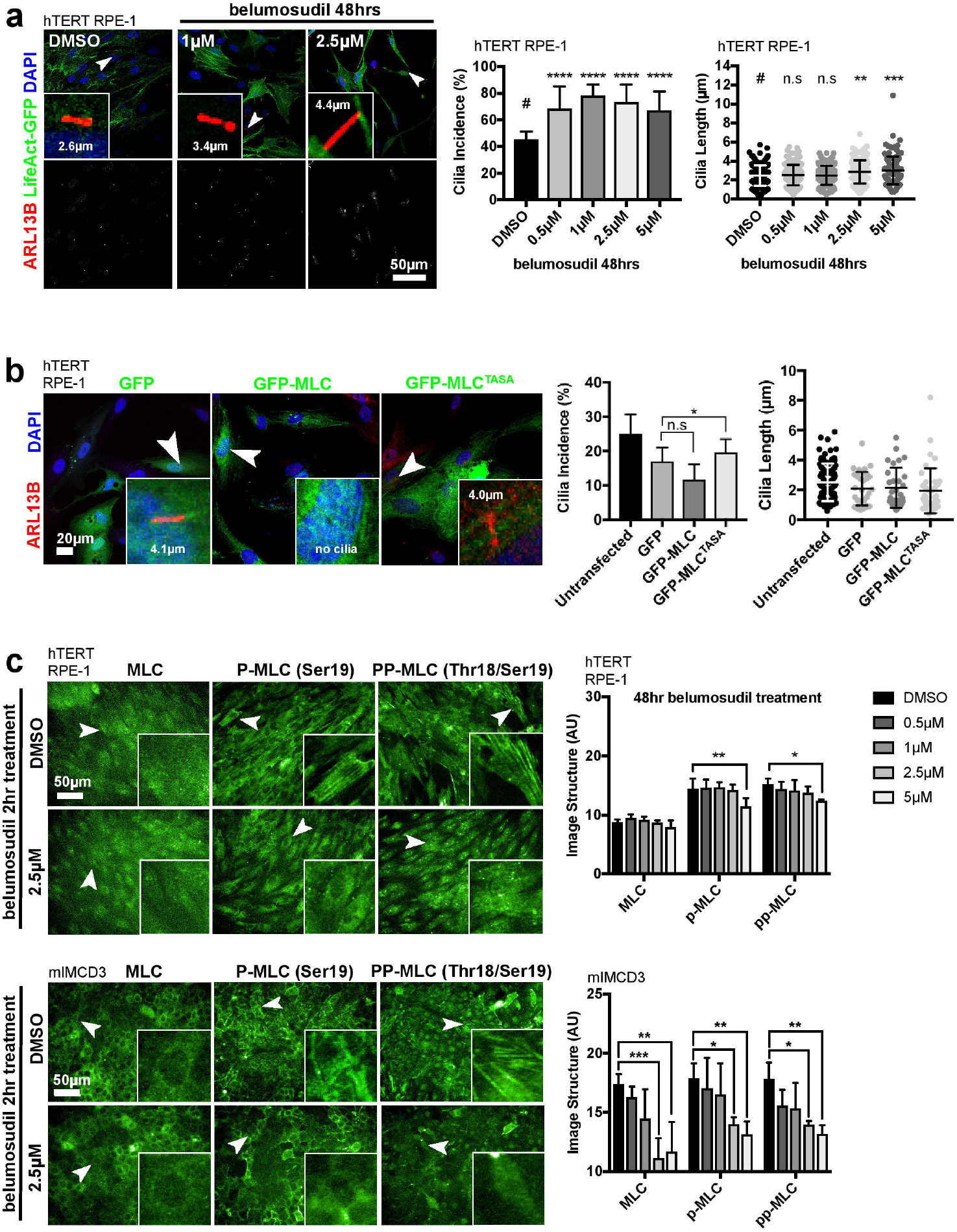
ROCK2 inhibition disrupts acto-myosin contraction and increases ciliogenesis. **(a)** Belumosudil treatment of hTERT RPE-1 cells for 48 hrs significantly increased cilia incidence across all concentrations (0.5 to 5 μM) and increased cilia length for concentrations ≥2.5 μM compared to vehicle (DMSO) control (n=3, minimum of 40 cilia measured per replicate). Mean cilia length for control treatment was 2.5 μm, compared to 2.9 μm and 3.0 μm at 2.5 μM and 5 μM belumosudil, respectively. Statistical significance calculated with one-way ANOVA with Dunnett’s test for multiple corrections (cilia incidence) and Kruskal-Wallis test with Dunn’s multiple comparisons test (cilia length) as indicated (* *p*<0.05, ** *p*<0.01, *** *p*<0.001, **** *p*<0.0001). **(b)** hTERT RPE-1 cells were transfected with GFP-tagged active MLC or constitutively inactive MLC^AA^ constructs (green) and stained for ARL13B (red). Arrowheads indicate cells displayed in magnified insets. Cells that overexpressed active MLC had a moderate but non-significant (n.s.) decrease in cilia incidence (p=0.0866), MLC^AA^ expression significantly increase cilia incidence compared to untagged GFP control (n=3). There were no significant changes in cilia length (n=3, p=0.6464, minimum of 25 cilia measured per condition). Statistical significance of pairwise comparisons is indicated (* p<0.05, unpaired two-tailed Student’s t-test). **(c)** Belumosudil inhibits the kinase activity of ROCK2 and changes non-muscle myosin IIA organization, visualized by myosin light chain (MLC) staining and the presence of MLC-associated acto-myosin structures in both hTERT RPE-1 (upper panels) and mIMCD3 (lower panels) cell-lines. Cells were treated with increasing concentrations of belumosudil for either 2 hrs or 48 hrs (n=3 for both time points). Antibodies marked MLC, p-MLC (mono-phosphorylated MLC) and pp-MLC (biphosphorylated MLC at Thr18 & Ser19), with representative immunofluorescence confocal microscopy images shown with 2.5 μM belumosudil for 2 hr treatments. p-MLC and pp-MLC visualize acto-myosin fibre-like structures in both hTERT RPE-1 (upper panels) and mIMCD3 (lower panels), visible in cells indicated by arrowheads and displayed in magnified insets. Automated high content image analysis of image texture quantified both p-MLC and pp-MLC fibre-like structures, with bar graphs showing significant decreases in both hTERT RPE-1 (top) and mIMCD3 (bottom) cells when treated with 5 μM belumosudil for 48 hrs. Statistical significance was calculated with a two-way ANOVA with Dunnett’s multiple comparisons test as indicated (* p<0.05, ** p<0.01, *** *p*<0.001). Scale bar = 50 μm.

### Acto-myosin contraction contributes to the regulation of ciliogenesis

ROCK2 has not been directly shown to be a negative regulator of ciliogenesis in previous studies, but downstream actin remodelling pathways activated by ROCK have provided indirect evidence for the importance of actin remodelling in controlling cilia incidence and length (27, 28). Thus, the current published evidence suggests that a highly complex network of pathways and regulators control the functional contribution of the actin cytoskeleton during ciliogenesis. ROCK2 is reported to regulate F-actin stability though direct phosphorylation of both LIMK2 and non-muscle myosin light chain 2 (MLC), the regulatory subunit of non-muscle myosin IIA, leading to activation of acto-myosin cell contractility (39). Local activation of apical acto-myosin contractility controls cell adhesion, polarity and morphogenesis (40). In particular, acto-myosin contractility is required for basal body migration to the apical cell surface (41) an essential mechanical event required prior to ciliogenesis, although the molecular details remain unclear.

To substantiate the contribution of ROCK2, we first tested the effect of chemical inhibitors of F-actin stabilisation (cytochalasin D) and non-muscle myosin II (blebbistatin). Consistent with a previous study (27), treatment with 0.5 μM cytochalasin D significantly increased cilia incidence over 16 hrs from 28.8 to 51.3%, and significantly increased cilia length in as little as 2 hrs by 11.7% (Supplementary Figure 6, A). Interestingly, 0.5 μM blebbistatin treatment also increased cilia incidence over 16 hrs from 25.3 to 44.7% (consistent with previous reports (41)) and 2.5 μM treatment significantly increased cilia length in just 2 hrs by 20.9% (Supplementary Figure 6, B). These data indicate that over the shorter time period of 2 hrs, treatment with either cytochalasin or blebbistatin caused significant increases in cilia length, whereas longer 16 hr treatments caused increases in cilia incidence. The increase in cilia incidence was more significant for treatment with blebbistatin, which inhibits acto-myosin contractions, than for cytochalasin D which inhibits F-actin stabilisation. This may indicate the relative contributions of both acto-myosin contractility and F-actin stabilisation to ciliogenesis, with a greater effect of actin polymerisation on cilia length over short time periods compared to a longer-term effect of acto-myosin contractility inhibition on cilia incidence.

We further validated the functional role of acto-myosin contractility during ciliogenesis by transfecting GFP-tagged wild-type rat myosin light chain (MLC) or a constitutively inactive non-phosphorylatable mutant of MLC (MLC^AA^) into hTERT RPE-1 cells (42). Expression of wild-type GFP-MLC did not significantly decrease cilia incidence or length, possibly because it was not readily phosphorylated by ROCK2. However, constitutively inactive GFP-MLC^AA^ did significantly increase cilia incidence compared to GFP only transfection control (Figure 6, C), phenocopying blebbistatin treatment and *Rock2* siRNA knock-downs. We therefore validated that siRNA knockdown and belumosudil both significantly reduced the presence of fibre-like structures associated with acto-myosin contractility. To verify that changes in actomyosin contractility were contributing to the cellular phenotypes observed in our previous experiments, we again used high content imaging to assess levels of phosphorylated nonmuscle myosin light chain 2 (MLC) and image structure analysis to quantify the levels of fibre-like structures associated with acto-myosin contractility. This confirmed that levels of phosphorylated MLC and acto-myosin contractility were reduced in experimental conditions that disrupted ROCK2 activity, specifically siRNA knock-down (Supplementary Figure 7, A and B) and belumosudil treatment (Figure 5, B).

## Discussion

In this study, two hypothesis-free approaches highlighted ROCK2 as a potential therapeutic target for the restoration of cilia in ciliopathy patients. The drug screen identified fasudil, a broadly selective ROCK inhibitor, as a chemical that can rescue cilia after knockdown of several known ciliopathy genes in different cellular disease models. Furthermore, increased cilia incidence with drug treatment was associated with rescue of ciliary-mediated Sonic Hedgehog signalling. The independent whole genome reverse genetics screen and subsequent validation experiments identified ROCK2 as the target protein that mediates a strong negative regulation of cilia incidence. Functional studies then verified that ROCK2 regulates ciliogenesis through regulation of acto-myosin contractility.

When activated by RhoA-GTP, both ROCK1 and ROCK2 phosphorylate several proteins and thus activate downstream pathways that regulate actin remodelling and dynamics. They have therefore been intensively studied to determine their roles in cell adhesion (43, 44), stress fibre and focal adhesion formation (45, 46), and cell motility/migration (reviewed in (47)). ROCK controls these cellular mechanisms by directly phosphorylating key actin regulators. ROCK phosphorylates LIMK2, which in turn phosphorylates cofilin to modulate Factin stabilisation, as well as MLC and MYPT1 which control acto-myosin contraction and the formation of stress fibres. ROCK also phosphorylates EZR regulating the linking of the actin cytoskeleton to the plasma membrane (48). Despite the overall sequence similarity of the two ROCK isozymes (65% overall, 93% in the kinase domain (49), our findings indicate that ROCK2 is the target molecule that mediates ciliogenesis rather than ROCK1. *ROCK1* knock-down does not phenocopy *ROCK2* knock-down, and residual ROCK1 activity cannot compensate for the chemical inhibition of ROCK2 kinase activity. These results agree with previous findings from knockout mice which showed that ROCK1 and ROCK2 have distinct roles and cannot compensate for one another (50, 51), although both ROCK1 and ROCK2 are ubiquitously expressed (33, 52, 53). ROCK1 and ROCK2 have been suggested to have different roles in the regulation of the cytoskeleton, with ROCK1 responsible for destabilising the actin cytoskeleton and regulating MLC phosphorylation, and ROCK2 thought to stabilise the actin cytoskeleton through regulation of cofilin phosphorylation (35). Regulation of cytoskeletal dynamics is required for both basal body docking and ciliary vesicle trafficking and is therefore an essential process of early ciliogenesis, axoneme elongation and maintenance (27, 34, 54, 55).

Fasudil was one of the first ROCK inhibitors identified (56) and is a class I ATP competitive kinase inhibitor of both ROCK isozymes (Supplemental Table 2). Previous drug repurposing and preclinical studies have now led to an open-label, single-centre Phase II clinical trial in China to investigate the efficacy and safety of fasudil for the neurodegenerative condition amyotrophic lateral sclerosis (NCT01935518). More recent laboratory and clinical studies have shown the efficacy of ROCK inhibitors in treating conditions such as asthma, cancer, erectile dysfunction, insulin resistance, chronic kidney disease, neuronal degeneration, osteoporosis, cerebral vasospasm, glaucoma, idiopathic pulmonary fibrosis, psoriasis, nonalcoholic steatohepatitis and graft versus host disease (reviewed in (57)). However, despite this, their good oral bioavailability and suitable half-lives, the compounds have only rarely entered licensed clinical use. Kinase profiling has indicated that the earlier first-generation ROCK inhibitors have significant off-target effects, likely because fasudil and hydroxyfasudil both target the ATP binding domain, which is 100% identical for both ROCK isozymes. The pharmacological effect of earlier ROCK inhibitors is also complicated by the potentially undesirable effects of ROCK1 inhibition (which include reduced blood pressure, increased heart rate and reversible reduction in lymphocyte counts) (57, 58). More recently, better, isozyme-specific inhibitors have been formulated, such as belumosudil (KD025) (59–61) and many others (62). Despite this, it is notable that these compounds are often missing from drug screening libraries and that many remain commercially unavailable. Lack of screening of specific ROCK1 or ROCK2 inhibitors might be masking inhibition of one or other isozyme as an effective target for the treatment of many other diseases.

Other screens to identify potential therapeutics for cystic kidney disease have also identified ROCK inhibitors. One study screened a library of 273 kinase inhibitors and identified thiazovivin (a pyrimidine-based non-selective ROCK inhibitor) as a significant hit (63). It is also interesting to note that the plant flavanoid, eupatilin, which was highlighted as a potential therapeutic for *CEP290*-related retinal dystrophy (11) and has been used to treat the *Rpgrip1l*^-/-^ mouse (64), has an indirect effect on actin cytoskeleton remodelling that is mediated by CEP290 (65). Several other screens have identified other biological targets for the restoration of cilia, for example a screen to identify drugs that restore cilia expression in cancer cells identified 118 compounds, many of which affect levels of cAMP, calcium or other ions (66). In agreement with our findings, ROCK inhibitors have also been shown to restore cilia in a Rho GTPase activating protein (GAP) mutant mouse model (*Arhgap35*^D34/D34^) that had reduced cilia incidence and length (67). They have also been identified as restoring cord /tubule formation whilst reducing cyst formation in a cystogenic mIMCD3 *Pkd1*^-/-^ 3D kidney model (68). In this study 155 kinase inhibitors were tested and 5 hit compounds were reported, of which all were ROCK inhibitors, including a selective ROCK2 inhibitor. This suggested that ROCK2, and not ROCK1, is the key target molecule, again in agreement with our findings.

Neither our drug screen nor reverse genetics screen assessed the effects of all screened chemicals or siRNA knock-downs on cilia length, since our high content imaging did not allow accurate analysis in mIMCD3 cells. However, another study showed that the non-selective ROCK inhibitor Y27632 caused an increase in cilia length due to reductions in actin stress fibre formation (69). Our validation experiments showed significant increases in cilia length in cells treated with si*ROCK2*, the ROCK2 specific inhibitor belumosudil, cytochalasin D, or with blebbistatin, dependant on length of treatment.

In conclusion, our work suggests that ROCK2 is a key mediator of cilium formation and function and have determined that ROCK2 inhibition has a broad effect in rescuing ciliary function over several ciliopathy disease classes. We propose that ROCK2 inhibition represents a novel, disease-modifying therapeutic approach for heterogeneous ciliopathies. We have identified fasudil as a compound that can restore cilia after knockdown and ROCK2 as the strongest negative regulator of ciliogenesis. Our data suggest that ciliogenesis is enhanced through two separate, but not mutually exclusive, effects on the actin cytoskeleton and acto-myosin contractions. These effects comprise a relatively quick increase in average cilia length (within 2 hrs), consistent with rapid inhibition of actin stabilisation in turn leading to increased ciliary vesicle transport. Secondly, a slower effect that increases overall cilia *incidence* (within 16 hrs), is consistent with longer term inhibition of acto-myosin contractility which likely caused enhanced basal body docking at the apical cell surface. Future work will focus on preclinical studies of fasudil or related isoquinolone derivatives as repurposable drug candidates. However, since fasudil is an off-patent, generic drug there are significant regulatory and commercial challenges to potential future repurposing of this candidate (12), newer more selective ROCK2 inhibitors may be more promising alternatives for ciliopathy disease indications. Indeed, belumosudil (known previously as KD-025) has been recently granted orphan drug status in the US, making it a potential candidate for drug repurposing.

## Methods

### Cell lines

Cell lines were sourced from American Type Culture Collection^®^ (ATCC^®^). Cells were used for screening between passage 15-25. mIMCD3 and hTERT RPE-1 mother celllines were previously verified using arrayCGH and RNA-sequencing (32) (Short Read Archive accession numbers SRX1411364, SRX1353143, SRX1411453 and SRX1411451). NIH3T3 “Shh-Light II” cells (PMID: 10984056) were a gift from Frédéric Charron, Montreal Clinical Research Institute. Cell-lines were maintained by weekly passaging under standard conditions and tested every 3 months for mycoplasma.

### Antibodies

The following antibodies were obtained commercially: anti-acetylated α-tubulin (clone 6-11B-1, Sigma T6793); anti-ARL13B (Proteintech 17711-1-AP); anti-ARL13B (clone N295B/66, NeuroMab) anti-myosin light chain II (CST 8505); anti-phosphorylated myosin light chain II (Ser19) (CST 3671); phosphorylated myosin light chain II (Thr18/Ser19) (CST 3674); anti-ROCK1 (clone C8F7, CST 4035); anti-ROCK2 (Bethyl A300-047A); anti-GAPDH (CST 2118); anti-β-actin (clone AC-15, Abcam Ab6276); anti-Smoothened homolog (Bioss antibodies, bs-2801R).

### Immunofluorescence imaging

Cells plated on slides or in 96-well plates were fixed using 4% paraformaldehyde or using ice-cold methanol followed by permeabilisation using PBS containing 0.1% Triton X-100. Cells were stained for immunofluorescence using standard methods. Cells in plates were stained using anti-ARL13B (1:6000) as a cilia marker. Cells on coverslips were stained for with ARL13B (1:2000) or acetylated α-tubulin (1:1000) as cilia markers. Slides were imaged as z-stacks using confocal microscopy (Nikon A1R Confocal Laser Scanning Microscope, controlled by the NIS-Elements C software). Confocal images were analysed for cilia length and incidence using FIJI image software (70) macros. Cilia and nuclei were highlighted as regions of interest (ROI) and were quantified for total number of each ROI, cilia length and total fluorescence using a custom written macro (Supplemental data 3). 96-well plates were stained for MLC, p-MLC (Ser19) and pp-MLC (Thr18/Ser19) (1:50) to image myosin light chain II and phosphorylated isoforms, and ARL13B (1:2000) to image cilia. Plates were imaged at 40X using a high-throughput microscope (PerkinElmer Operetta^®^ microscope controlled using Harmony^®^ high-content analysis software). Image data was analysed using Columbus^™^ Image Data Storage and Analysis system (PerkinElmer). Image structure of MLC staining was analysed using the image structure analysis block, with Gabor image structure analysis (scale: 2px, wavelength: 8, number of angles: 8 with regional intensity).

#### Small molecule library and other compounds

The Tocriscreen Total drug screen library of 1120 biologically active compounds with wide chemical diversity was obtained from Tocris (product no. 2884). Compounds were supplied at 10 mM in DMSO. Fasudil hydrochloride was obtained from Cell Guidance Systems (SM-49), belumosudil (KD-025) from Cayman Chemical (17055), and hydroxyfasudil (S8208) and ripasudil (S7995) were obtained from Selleckchem.

#### Drug Screening: Primary and secondary screen

mIMCD3 cells were seeded in 96-well ViewPlates (PerkinElmer) at 8000 cells per well in Opti-MEM. The cells were reverse transfected with si*Rpgrip1l* or negative control siScr using Lipofectamine RNAiMAX as detailed previously. Control transfections of si*Ift88, siMks1* and a mock transfection were also included. Chemicals were added 24 hrs after cell plating at a final concentration of 10 μM. The final concentration of DMSO vehicle in each well was 0.1%. Cells were fixed and stained 72 hrs after seeding and 48 hrs after addition of chemicals. Cells were fixed using ice-cold methanol and permeabilised using 0.1% Triton X-100 in PBS. Cells were stained using anti-ARL13B (Proteintech) at 1:6000, followed by staining with goat anti-rabbit Alexa Fluor 488, (A11034, Fisher) at 1:2000, DAPI (Fisher) at 1:5000 and TOTO-3 (Fisher) at 1:5000. High content imaging and analysis was as described above. For the primary screen, robust *z* scores were normalized to account for variation between plates. Mean values for two passages were used to identify chemicals that significantly restored cilia after knockdown without significant effects on cell number.

#### Drug screening: Tertiary screen

hTERT RPE-1 cells were seeded at 20,000 cells per well in DMEM/F12 0.2% FCS in 96-well ViewPlates coated with Matrigel Matrix (Corning). Similar methodology was used to the secondary screen except that cells were reverse transfected with si*IFT88* as well as si*RPGRIP1L* and siScr.

#### MTT assay

hTERT RPE-1 cells were seeded at 11,200 per well in DMEM/F12 0.2% FCS in a 96-well plate. mIMCD3 cells were seeded at 12,800 cells per well in Opti-MEM in a 96-well plate. Fasudil hydrochloride was diluted in DMSO and added to cells at a final concentration of 0.1, 0.3, 1, 3, 10, 30 and 100 μM. At each time-point, media was removed and 50 μl of MTT reagent at 1 mg/ml added. Plates were incubated for 3 hours in darkness at 37°C. Any remaining MTT solution was removed and dark blue formazan precipitates were solubilised in 100 μl propan-1-ol. Optical density was measured at 570 nm using a Mithras Berthold LB940 plate reader.

#### Generation of CRISPR/Cas9 heterozygote cell lines

hTERT RPE-1 cells were CRISPR edited using guide RNAs (gRNAs) designed using the online CRISPR Design tool (crispr.mit.edu [no longer available]) to target the coding exons of *RPGRIP1L* (GGTCCAACTGATGAGACTGC) and *IFT88* (GGGCCCCTTGACTGACTAAG). gRNA for *TMEM67* (TAACAAATGTTGGCTCACAT) was designed using Benchling (www.benchling.com/). *BbsI* restriction sites were added and reverse HPLC-purified oligonucleotides were obtained commercially. Oligos were annealed and then cloned into the pX458 CRISPR/Cas9 expression vector. The cloned vectors were transfected into hTERT RPE-1 cells using Lipofectamine 2000. Transfected cells were single cell sorted by FACS into 96-well plates 24 hours post transfection. Genomic DNA was extracted from confluent colonies, the targeted regions amplified and colonies with heterozygous variants identified using a T7 endonuclease digestion. Variants were identified using Sanger sequencing.

#### Dose response assays

hTERT RPE-1, hTERT RPE-1 *IFT88*^+/-^, hTERT RPE-1 *RPGRIP1L*^+/-^, hTERT RPE-1 *TMEM67*^+/-^ and hTERT RPE-1 *TMEM67*^-/-^ cell lines were seeded at 9000 cells per well in DMEM/F12 plus 0.2% FCS in 96-well ViewPlates coated with Matrigel matrix. mIMCD3 cells were seeded at 9000 cells per well in Opti-MEM in uncoated 96-well View plates. Fasudil hydrochloride, hydroxyfasudil and ripasudil were diluted in DMSO. Chemicals were added and cells were stained as previously described.

#### Luciferase Assay

NIH3T3 “Shh-Light II” cells were reverse transfected in Opti-MEM and seeded at 14,400 per well in 96-well plates. Fasudil hydrochloride, Smoothened agonist (SAG; Cayman 11914) or vehicle controls were added after 24 hours to final concentrations of 10 μM and 250 nM respectively. Cells were lysed using 30 μl passive lysis buffer (Promega). 10 μl of cell lysate was analysed using a Mithras Berthold LB940 plate reader in a white 96-well plate (Greiner) using injectors to dispense 20 μl luciferase assay reagent (Promega) followed by 20 μl Stop n Glo substrate (Promega) diluted 1:1 with filtered dH_2_O. Values were expressed as ratios of firefly luciferase: *Renilla* luciferase normalised to the values for the unstimulated DMSO vehicle control treated cells.

#### SMO occupancy assay

mIMCD3 cells were seeded on coverslips at 1.1 x 10^5^ cells in 1ml Opti-MEM. Chemicals were added after 24 hrs and cells were fixed and stained after 72 hrs using 1:300 anti-SMO, 1:2000 anti-ARL13B (Neuromab) antibodies followed by goat anti-rabbit 488 and goat antimouse 568 both at 1:2000 and DAPI at 1:500. All antibodies were diluted in 3% goat serum. Slides were imaged using confocal microscopy as described above. Images were analysed using Columbus to identify cilia as regions of interest and the SMO occupancy within those regions. Values of total SMO area: cilia number ratios were used to control for variable cilia size.

### Analysis of Primary whole genome siRNA screen data

Primary whole genome screen data was published in Wheway, 2015 (32). The secondary screen hit list was generated by filtering out hits that had any significant changes to cell number in any experimental replicates (robust z score for cell number 2 ≥ x ≥ −2), significantly increase cilia incidence in both experimental replicates (robust z score for cilia incidence x ≥ 2). Final hits were selected based on siRNA specificity, and existence of human orthologues.

### Secondary siRNA screen

Secondary screening was performed following the previously published whole genome screen protocol (32). 2.5 μl of ON-Target Plus siRNA libraries (Dharmacon) were reverse transfected into mIMCD3 cells (8000 cells/well) as 2 μM siRNA (final concentration 50 nM) using Lipofectamine RNAiMAX in Opti-MEM. Each 96-well plate included 8 different controls duplicated in columns 1 and 12. si*Plk1* as a control for transfection; *siMks1, siIft88* and *siRpgrip1l* are positive controls for cilia loss; negative controls were siScr, si*MLNR* (designed against a human gene with no target in the mouse genome), and mock transfection with transfection reagent only. siRNA transfections were incubated for 72 hrs. Plates were fixed in ice cold methanol and stained high-throughput using fluid dispensers. Plates were immuno-stained for acetylated α-tubulin (1:4000) and stained with DAPI (1:2000) and TOTO-3 (1:4000). Plates were imaged at 20x using a high-throughput microscope (PerkinElmer Operetta^®^ microscope controlled using Harmony^®^ high-content analysis software). Image data was analysed using Columbus^™^ Image Data Storage and Analysis system (PerkinElmer). Image data was analysed to identify whole nuclei, cell boundaries and cilia. Robust *Z* scores were used to determine significant hits within each experimental replicate. Negative controls were pooled (siScr, *siMNLR*, mock transfection) within each experimental replicate and compared to individual siRNA knockdown values for each quantified phenotype. Average robust *Z* scores were calculated, and hits were classed as having average robust *z* score for cilia incidence (*z*_cilia_) ≥ 2.

### Western Blotting

Whole cell lysates were separated by SDS-PAGE and blotted onto PVDF membranes using the NuPAGE system and reagents (Invitrogen^™^). Membranes were blocked in 5% [w/v] milk/TBST and then immunoblotted for ROCK2 (A300-046A, 1:1000), ROCK1 (#4035, 1:0000), IFT88 (Proteintech 13967-1-AP, 1:1000), RPGRIP1L (Proteintech 55160-1-AP, 1:1000). GAPDH (Cell Signalling Technology #2118, 1:5000), vinculin (Sigma V4505, 1:5000) or β-actin (Abcam Ab6276, 1:10000) antibodies were used as loading controls.

### Plasmids

GFP-mROCK2 was a gift from Alpha Yap (Addgene plasmid # 101296). 500 ng of plasmid DNA was transfected into 1×10^5^ hTERT RPE-1 cells in DMEM/F12 medium containing 10% fetal calf serum (FCS), on 13 mm coverslips using Lipofectamine2000 (1:3 ratio DNA:Lipofectamine). Transfection complexes were removed after 4 hr and replaced with fresh medium. At 24 hrs post transfection, cells were serum-starved in DMEM/F12 0.2% FCS for 24 hrs before coverslips were fixed and prepared for immunofluorescence imaging.

### Inhibitor treatments

1×10^5^ cells were grown on 13mm round coverslips in DMEM/F12 containing 10% FCS for 24 hrs and then serum-starved as above. Cells were then treated with either vehicle control (DMSO) or chemical inhibitors at varying concentrations, diluted to final working concentrations in DMEM/F12 0.2% FCS. Coverslips were fixed and prepared for immunofluorescence imaging.

#### Statistical analysis and screen quality controls

Robust *z* scores (13) for cell number (*z*_cell_) and the percentage of cells with a single cilium (*z*cilia) were calculated for all results, compared to the median and median absolute deviation (MAD) of all positive and negative controls in a processed batch. This allowed application of meaningful statistical cut-off values (*z*_cilia cutoff_ and *z*_cell cutoff_) for the identification of significant “hits” affecting ciliogenesis based on the median and MAD of positive and negative controls per four-plate batch (14). The strictly standardised mean difference (SSMD) of the percentage of cells with a single cilium in *Ift88* and *Rpgrip1l* knockdowns, compared to all negative controls was consistently >1.5 reflecting the suitability of these siRNAs as positive controls for effects on ciliogenesis, and the consistency and quality of the screen (14, 15). For cell number, cilia incidence and cilia length measurements normal distribution of data was confirmed using the Kolmogorov-Smirnov test (GraphPad Prism software). Pair-wise comparisons were analysed with Student’s two-tailed t-test or one-way analysis of variance (ANOVA) using InStat (GraphPad Software). Significance of pair-wise comparisons indicated as: ns not significant, * *p*<0.05, ** p<0.01, *** p<0.001, **** p<0.0001. Results reported are from at least three independent experimental replicates, apart from drug and reverse genetics screens which comprised of two replicates (runs). Error bars on graphs represent standard error of the mean (s.e.m.).

## Author contributions

AVRL and CELS designed and conducted experiments, acquired and analysed data and wrote the manuscript and are therefore designated as equal first authors. SN conducted experiments, acquired and analysed data. RT provided reagents. JB and RF designed the research study and provided reagents. CAJ designed the research study, analysed data and wrote the manuscript.

## Acknowledgments

CELS, JB, RF and CAJ was funded by Action Medical Research (project grant no. GN2628), SN and CAJ were funded by NewLife the Charity for Disabled Children (grant no. 12-13/11). AVRL and CAJ were funded by UKRI (BBSRC-SFI joint project grant no. BB/P007791/1). Funding to pay the open access charges was provided by UKRI.

