## Supplemental Data 3 for "Drug and siRNA screens identify ROCK2 as a therapeutic target for ciliopathies"

run("Z Project...", "projection=[Max Intensity]");
//run("Channels Tool...");
title = getTitle();
run("Split Channels");
selectWindow("C4-" + title);
close();
selectImage("C3-" + title);
run("Subtract Background...", "rolling=5"); // removes background. If this is too harsh increase the "rolling" value.
run("Median...", "radius=3"); 
//run("Gamma...", "value=0.70"); //optional line may help to include for some noisy images.
setAutoThreshold("Yen dark");
setOption("BlackBackground", false);
run("Convert to Mask"); // Generate binary image.
run("Dilate");
run("Dilate"); // makes each thresholded cilia one pixel wider, to join any gaps. Possible to omit if results show cilia are too long.
run("Skeletonize");
run("Set Measurements...", "area perimeter redirect=None decimal=3");
run("Analyze Particles...", "size=0.1-100 show=[Overlay Masks] display exclude clear include summarize add in_situ");
        for(n=0; n<nResults;n++) {
	        setResult("Cilia Length", n, ((getResult("Perim.", n))/2));
        } // this for loop takes the results of each cilia perimeter, divides by 2 to give the length and adds it to the results table.
selectWindow("C1-" + title);
run("8-bit");
run("Subtract Background...", "rolling=50");
run("Auto Threshold", "method=Huang");
run("Make Binary");
run("Watershed");
run("Set Scale...", "distance=0 known=0 pixel=1 unit=pixel");
run("Analyze Particles...", "size=L-H pixel circularity=0.70-1.00 show=Nothing display exclude clear summarize");
selectWindow("Summary");
selectWindow("Results");
selectWindow("C3-" + title);
close();
