## Supplemental Table for "Drug and siRNA screens identify ROCK2 as a therapeutic target for ciliopathies"

**Supplemental Tables**

| Compound | $IC_{50}$ ROCK1 (µM) | $IC_{50}$ ROCK2 (µM) | Details | Source |
|---|---|---|---|---|
| Fasudil | 0.3 | "similar" to 0.3 | 1 µM ATP, human ROCK1 (aa 3-543) expressed in a baculovirus system (1) | (2) |
| | N/A | 1.9 | 100 µM ATP, human ROCK2, expressed in a baculovirus system | (3) |
| | 1.2 | 0.82 | No details | (4) |
| | 10.7 ± 2 | | 250 µM ATP, bovine brain derived ROCK[A] | (5) |
| | 2.670 ± 0.270 | 1.290 ± 0.080 | 40 µM ATP, human ROCK1 and ROCK2[B] | (6) |
| | 0.844 (1.174-0.515) | 0.691 (0.893-0.489) | 10 µM ATP, human ROCK1 (aa 1-535), human ROCK2 (aa 1-552) | (7) |
| | N/A | 0.158 | No details | (8)[C] |
| Hydroxyfasudil | 0.73 | 0.72 | No details | (4) |
| Ripasudil | 0.051 | 0.019 | 1 µM ATP, human ROCK1 (aa 1-477), ROCK2 (aa 1-553) expressed in a baculovirus system | (9) |
| KD025 | 24 | 0.105 | 10 µM ATP, human ROCK1 (aa 1-535), human ROCK2 (aa 1-552) expressed in a baculovirus system | (10) |
| | >10 | 0.059 | 10 µM ATP, human ROCK1 (aa 1-535), human ROCK2 (aa 1-552) expressed in a baculovirus system | (11) |
| | 5.1 | 0.07 | No details | Keystone Symposia on Molecular and Cellular Biology[D] |
| | 9.8 | 0.16 | No details | Paris NASH meeting[E] |

**Supplemental Table 1: $IC_{50}$ values for the ROCK inhibitors used in this study.** Note that the ATP concentration in cells is in the mM range, which is higher than that used for in vitro assays. However, in vivo activity is likely to be significantly lower than in vitro activity due to competition for ATP. Cell based assay $IC_{50}$ values are often 100-1000 fold higher than in vitro equivalents (12). Alternative names: fasudil (hydrochloride) / HA1077, hydroxyfasudil / HA1100, ripasudil / K115, KD025 / SLx-2119 / belumosudil.

[A] It is unclear whether unpurified ROCK, ROCK1 or ROCK2 was assayed, but many manufacturers specify this $IC_{50}$ value for ROCK1.

[B] The authors highlight differences in ATP $K_m$ values for ROCK1 and ROCK2, concluding that fasudil is not selective for either isoform.

[C] This reference is a review article with no experimental reference specified for the information stated.

[D]Keystone symposia on Molecular and Cellular Biology, January 2019: https://www.redxpharma.com/app/uploads/2019/01/ROCK2-poster-NASH-Keystone_Jan-2019_Emily-Offer.pdf

[E]Paris NASH meeting, July 2019:
https://www.redxpharma.com/app/uploads/2019/07/ROCK2-poster-NASH-July-2019_Emily-Offer.pdf

| Compound | $K_i$ ROCK1 µM | $K_i$ ROCK2 µM | $K_i$ ROCK µM | Details | Source |
|---|---|---|---|---|---|
| Fasudil | | | 0.35 | 100 µM ATP, bovine brain derived ROCK | (13) |
| | 0.145 ± 0.007 | 0.112 ± 0.008 | | 10 µM ATP, human ROCK1 (aa 1-535), human ROCK2 (aa 1-552) expressed in a baculovirus system | (11) |
| | | | 0.4 | 100 µM ATP, kinase domain of ROCK (aa 6-553) | (14) |
| | | 0.271 ± 0.014 | | 40 µM ATP, human ROCK2 | (6) |
| Hydroxyfasudil | | | 0.56 | No details | (13) |
| KD025 | >10 | 0.041 ± 0.002 | | 10 µM ATP, human ROCK1 (aa 1-535), human ROCK2 (aa 1-552) | (11) |

**Supplemental Table 2: $K_i$ values for the ROCK inhibitors included in this study.** Note that $K_i$ values for ripasudil are unavailable (15).

Alternative names: fasudil (hydrochloride) / HA1077, hydroxyfasudil / HA1100, ripasudil / K115, KD025 / SLx-2119 / belumosudil.

[A] Ito et al. (13) cites Seto et al. (16) but there is no information present for hydroxyfasudil
