## Supplemental Figure for "Drug and siRNA screens identify ROCK2 as a therapeutic target for ciliopathies"

**Supplemental Figures**

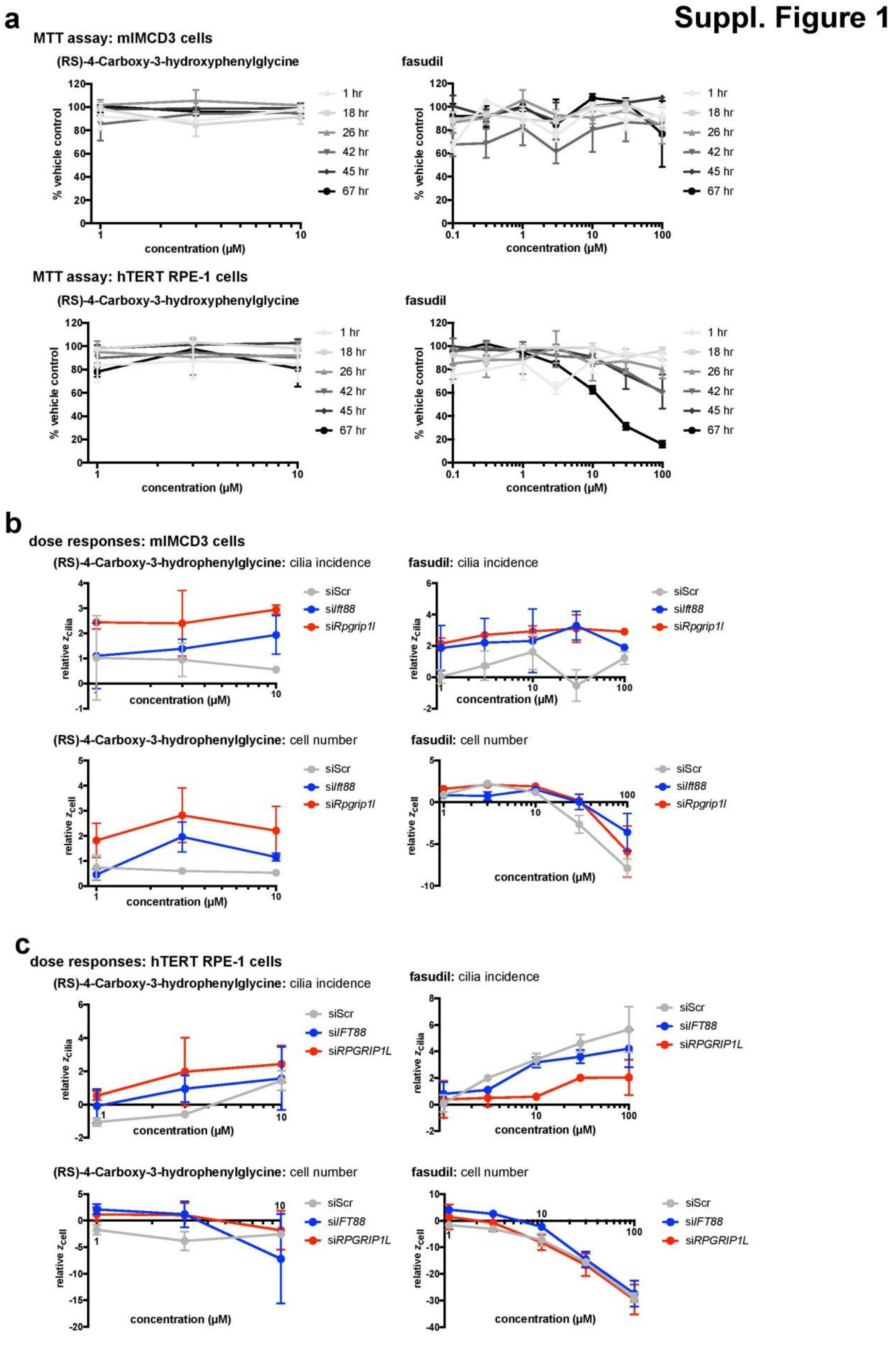


**Supplemental Figure 1: Fasudil dose response and cell viability assays in ciliopathy gene knockdown hTERT RPE-1 cells**.

**(a)** Dose response MTT assays of cell viability in mIMCD3 (upper two panels) or hTERT RPE-1 cells (lower two panels) treated with either (RS)-4-carboxy-3-hydrophenylglycine (left) or fasudil (right). Values are normalised to vehicle-treated control cells at each timepoint (1 to 67 hr treatments) for n=2 experimental replicates, each with n=3 technical replicates. Error bars represent range.

**(b)** Dose response assays in mIMCD3 cells with bar graphs showing $z_{cilia}$ (top panels) and $z_{cell}$ (bottom panels) values relative to vehicle (DMSO) treated cells for either (RS)-4-carboxy-3-hydrophenylglycine (left) or fasudil (right). Concentration range for (RS)-4-carboxy-3-hydrophenylglycine was 1 to 10 µM due to limited solubility in DMSO; fasudil was tested 1 to 100 µM. Cells were reverse transfected with siScr control (grey), si*Ift88* (blue) or si*Rpgrip1l* (red), as indicated. Values are for n=2 experimental replicates, each with n=3 technical replicates. Error bars present range. Significant increases / decreases are denoted by values > 2 / < -2

**(c)** Dose response assays in hTERT RPE-1 cells with bar graphs showing $z_{cilia}$ (top panels) and $z_{cell}$ (bottom panels) values relative to vehicle (DMSO) treated cells for either (RS)-4-carboxy-3-hydrophenylglycine (left) or fasudil (right). Concentration range for (RS)-4-carboxy-3-hydrophenylglycine was 1 to 10 µM due to limited solubility in DMSO; fasudil was tested 1 to 100 µM. Cells were reverse transfected with siScr control (grey), si*IFT88* (blue) or si*RPGRIP1L* (red), as indicated. Values are for n=2 experimental replicates, each with n=3 technical replicates. Error bars present range. Significant increases / decreases are denoted by values > 2 / < -2

**Suppl. Figure 2**

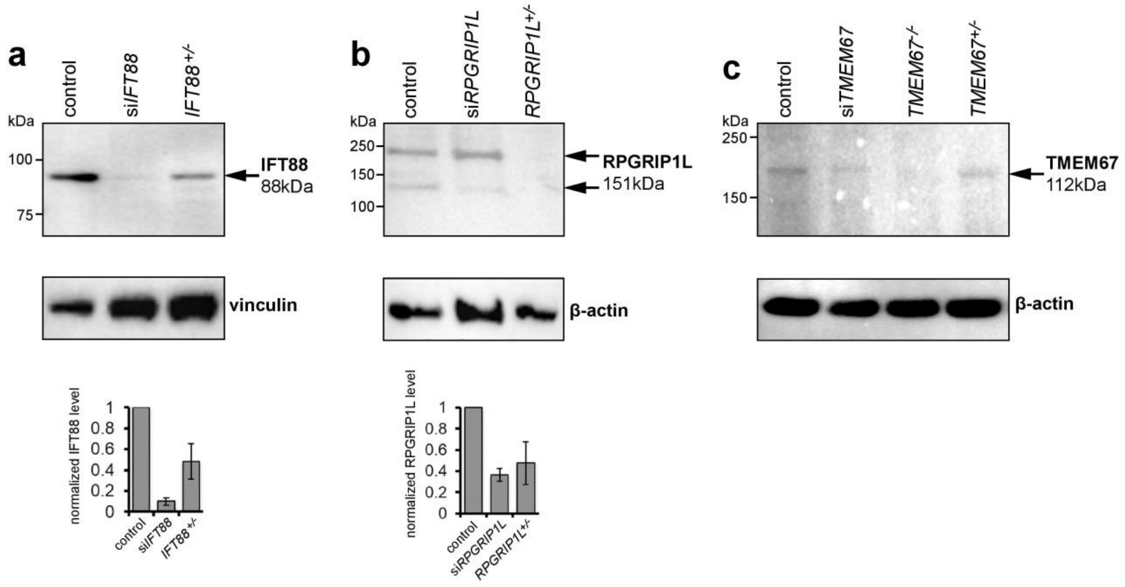


**Supplemental Figure 2: Assessment of relevant protein levels in CRISPR Cas9 edited cell lines.**

(a) Western blot showing the relative IFT88 protein levels in hTERT RPE-1 cells transfected with siScr or si*IFT88* or CRISPR-Cas9 engineered heterozygous knockout *IFT88* ((NM_175605.5):[c.371_408delinsAAGAAAAAAG,p.(P124Qfs*15)]) hTERT RPE-1 cells. Protein levels were quantified using densitometry relative to vinculin (n=3 experimental replicates). An arrow indicates the bands used for quantification, predicted weight 88 kDa.

(b) Western blot showing the relative RPGRIP1L protein levels in hTERT RPE-1 cells transfected with siScr (control) or si*RPGRIP1L* or CRISPR-Cas9 engineered heterozygous knockout *RPGRIP1L* ((NM_015272.5):[c.15_37del, p.(D6Cfs*35)]) hTERT RPE-1 cells. Protein levels were quantified using densitometry relative to β-actin (n=3 experimental replicates). Arrows indicate the bands that represent the two main protein isoforms detected, the predicted weight for the protein detected by this antibody was stated as 151 kDa but neither of the bands detected were of this size. The lower bands were used for quantification.

(c) Western blot showing the relative TMEM67 protein levels in hTERT RPE-1 cells of cells transfected with siScr (control) or si*TMEM67* or CRISPR-Cas9 engineered heterozygous knockout *TMEM67* ((NM_153704.6): [c.369delC, p.(E124Kfs*12)]) and homozygous knockout *TMEM67* ((NM_153704.6):[c.519delT, p.(C173Wfs*20)];[c.519dupT, p.(E174*)]) hTERT RPE-1 cells. The bands detected were of a larger weight than was expected (predicted weight detected by antibody was 112 kDa), possibly due to post translational modifications. High content imaging indicated a reduction in cilia incidence compared with the WT parent line for the CRISPR-Cas9 engineered *TMEM67*$^{+/-}$ and the *TMEM67*$^{-/-}$ lines of 59% and 89% respectively (data not shown).

#### Suppl. Figure 3

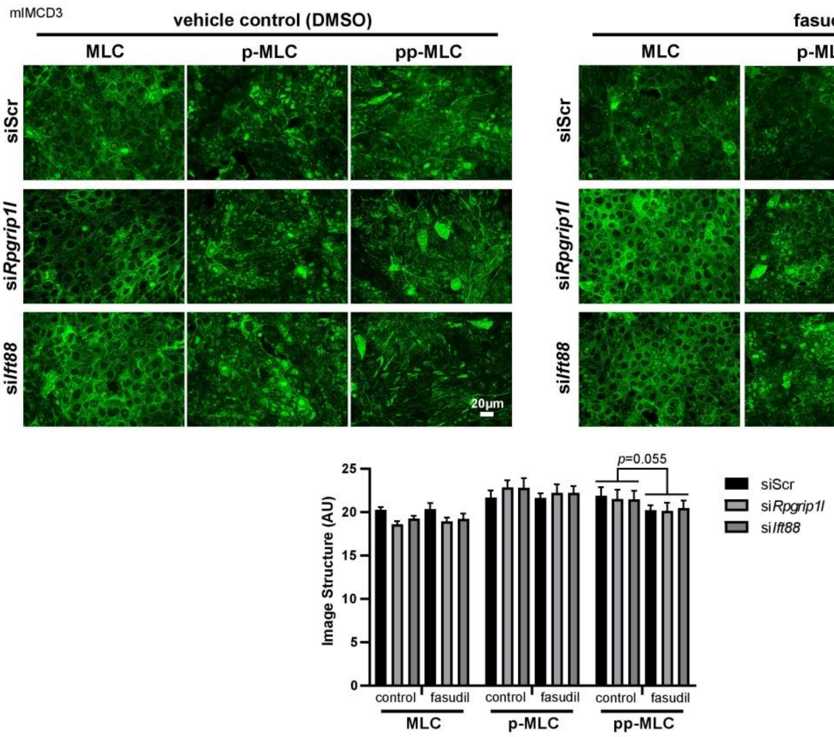


**Supplemental Figure 3: Fasudil does not significantly affect acto-myosin contractility.**

High content imaging of mIMCD3 cells treated with siScr, si*Rpgrip1l* or si*Ift88* and either DMSO (left panel) or 10 µM fasudil (right panel) to show levels of changes non-muscle myosin IIA organization. Acto-myosin structures were visualized by myosin light chain (MLC) staining and the presence of MLC-associated acto-myosin structures. Antibodies marked MLC, p-MLC (mono-phosphorylated MLC) and pp-MLC (biphosphorylated MLC at Thr18 & Ser19), with representative immunofluorescence confocal microscopy images shown. Automated high content image analysis of image texture quantified MLC, p-MLC and pp-MLC fibre-like structures, with bar graphs showing decreases in texture for pp-MLC that were not significant when cells were treated with 10 µM fasudil for 48 hrs. Statistical significance was calculated with a two way, unpaired Student's t-test as indicated (n=3 experimental replicates, each with n=4 technical replicates). Scale bar = 20 µm.

**Suppl. Figure 4**

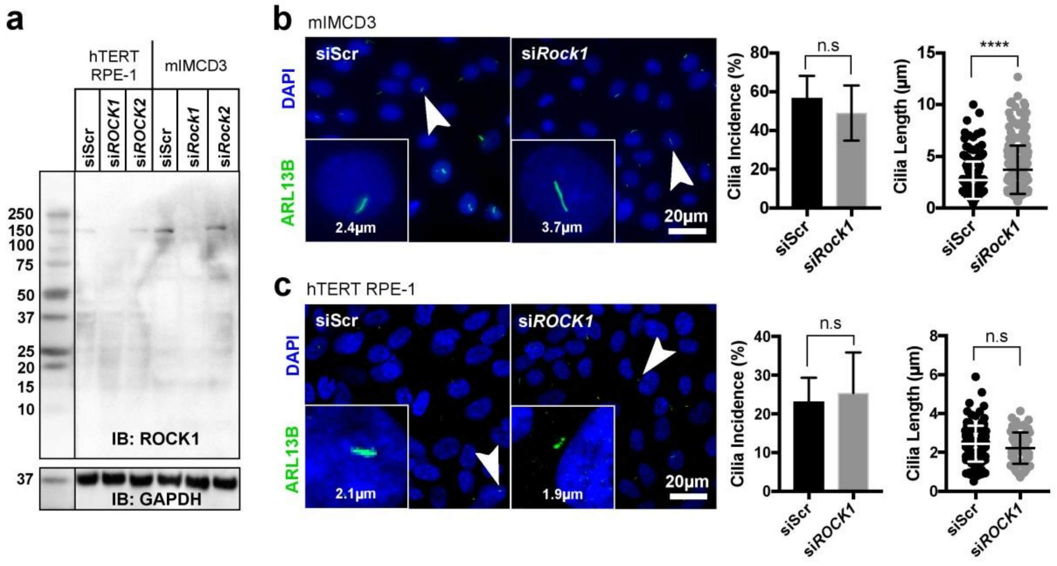


**Supplemental Figure 4: ROCK1 does not significantly modulate ciliogenesis.**

**(a)** Specificity of *ROCK1* siRNAs demonstrated by western blots following siRNAs knockdowns of either human *ROCK1* or mouse *Rock1* (expected molecular weight 158 kDa). GAPDH is the loading control.

**(b)** Representative confocal microscopy images of mIMCD3 cells knocked down with si*Rock1* or siScr negative control and stained for cilia marker ARL13B (green). Representative examples of cilia length measurement are shown for each condition. Bar graphs show a slight decrease in cilia incidence following si*Rock1* knockdown (n=3, $p$=0.3724) but a significant increase in mean cilia length from 2.7 µM to 3.7 µM (**** $p$<0.0001, n=3, at least 50 cilia measured per experimental replicate).

**(c)** Representative confocal microscopy images of hTERT RPE-1 cells treated with si*ROCK1* or siScr negative control as for (b) Bar graphs show a moderate but not significant increase in cilia incidence following si*ROCK1* knockdown (n=3, $p$=0.0891). Mean cilia length was decreased following si*ROCK1* knockdown, from 2.4 µM to 2.2 µM (n=3, $p$=0.1711, at least 50 cilia measured per experimental replicate). Statistical significance of pairwise comparisons was calculated with Student's two-tailed t-test (for cilia incidence) and the Mann-Whitney U-test (for cilia length).

#### Suppl. Figure 5

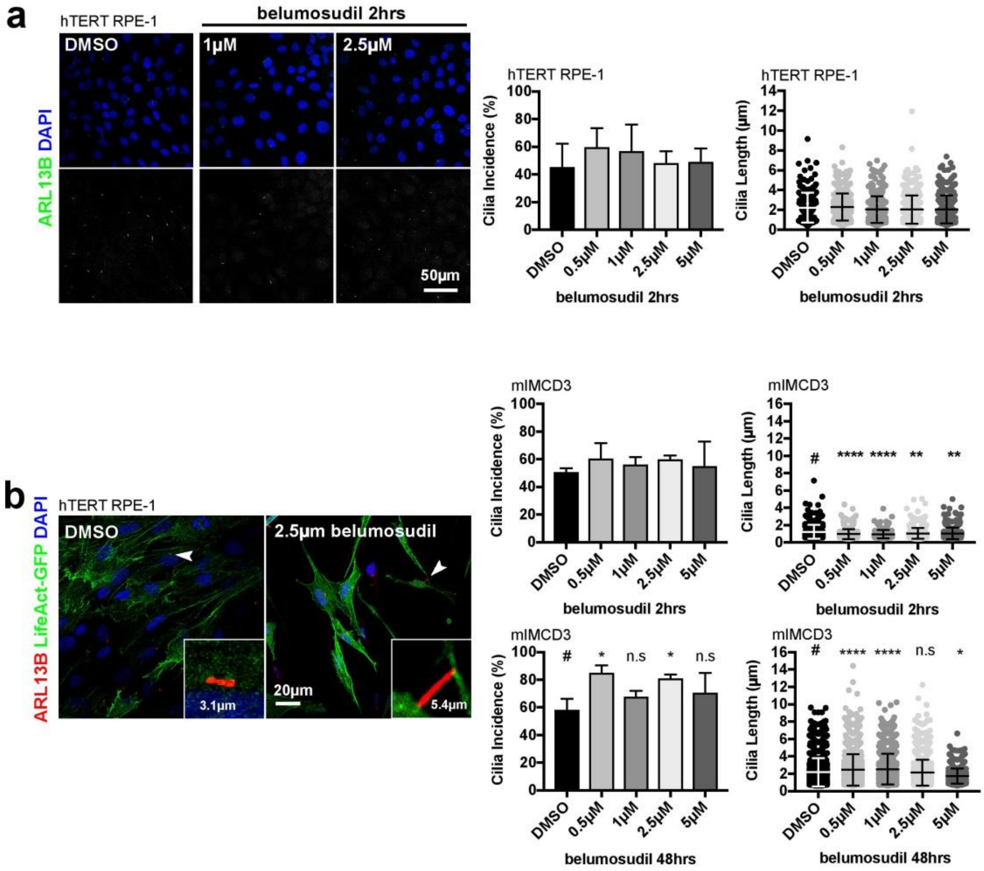


**Supplemental Figure 5: Treatment with belumosudil for 48 hrs significantly increases cilia incidence and length.**

**(a)** 2 hr treatment with belumosudil after 24 hrs serum starvation did not cause significant changes to either cilia incidence in hTERT RPE-1 (left panel) or mIMCD3 (right panel) cells. Data was confirmed to be normally distributed using D'Agostino-Pearson omnibus K2 test. Significance was then calculated using one-way ANOVAs with Dunnett's test for multiple corrections. Error bars represent S.D.

**(b)** 48 hr treatment of with belumosudil in serum starvation conditions led to significant increases in cilia incidence (left panels) and length (right panels) in both hTERT RPE-1 (upper panels) and mIMCD3 (lower panels) cells compared to DMSO-treated control cells (n=3 experimental replicates, with at least 100 cilia measured per experimental replicate). However, there was also a clear reduction in cell number and changes in cell morphology over this time period (images). Data for cilia incidence was confirmed to be normally distributed using D'Agostino-Pearson omnibus K2 test, whereas cilia length data was not normally distributed. Significance was then calculated using one-way ANOVAs with Dunnett's test for multiple corrections (cilia incidence) and Kruskal-Wallis tests with Dunn's multiple comparisons test (cilia length). *=p<0.05, **=p<0.01, ***=p<0.001, ****=p< 0.0001. # is the control all data sets were compared to. Error bars represent S.D.

### Suppl. Figure 6

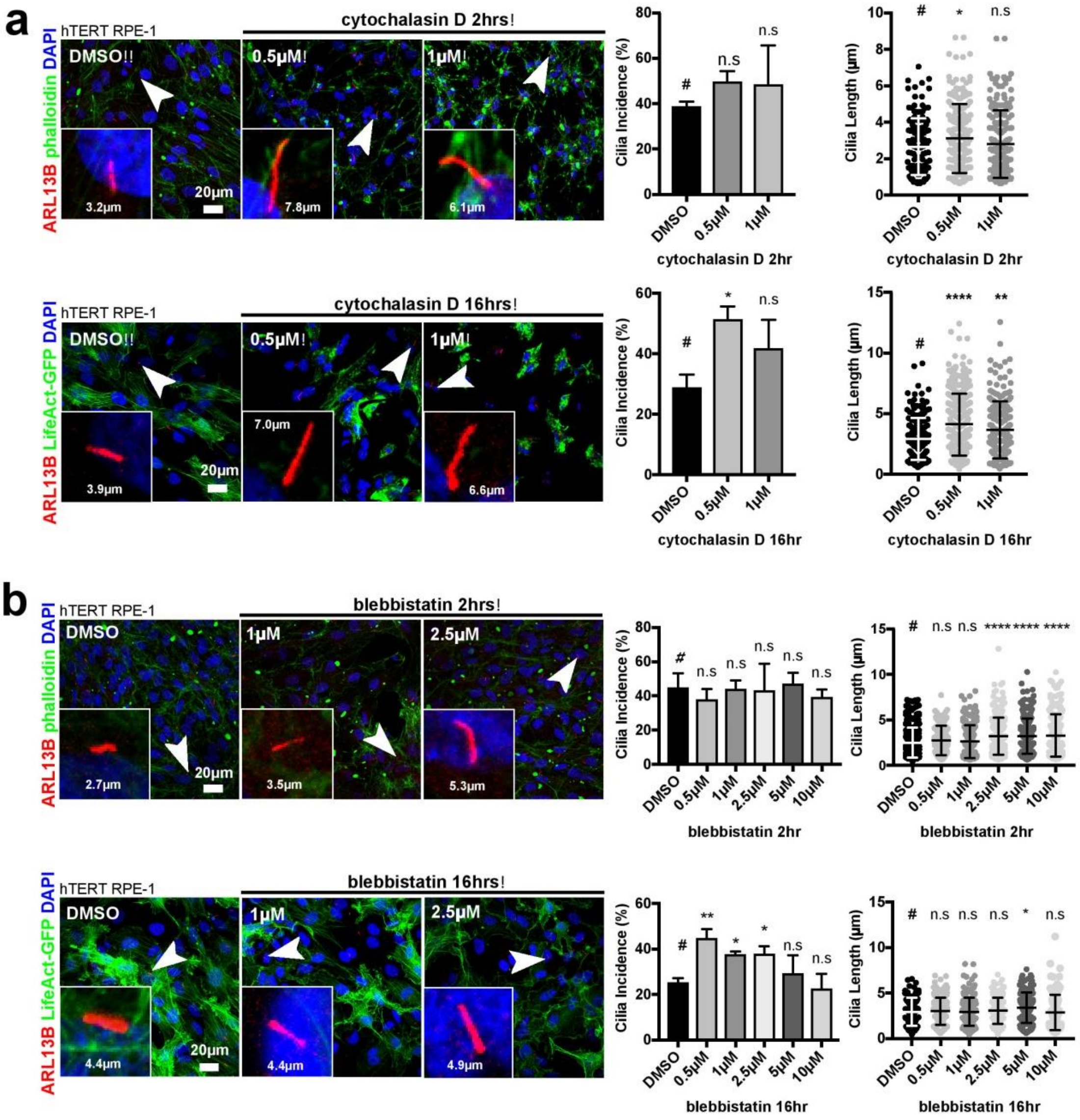


**Supplemental Figure 6: Treatment with cytochalasin D and blebbistatin increases ciliogenesis.**

**(a)** Pharmacological inhibition of F-actin stabilisation though treatment with cytochalasin D increased ciliogenesis. hTERT RPE-1 cells stably transfected with LifeAct-GFP (green) were treated for either 2 hrs (upper panels) with cytochalasin D or vehicle control after 24 hours of serum starvation, or for 16 hrs (lower panels) with cytochalasin D or vehicle control in serum starvation media (all conditions n=3). Representative confocal microscopy images show cells stained with cilia marker ARL13B (red) with arrowheads indicating cilia displayed in magnified insets and their measured lengths. Bar graphs quantitate the effect of inhibition with cytochalasin D at the indicated concentrations on cilia incidence and length. Cells treated with cytochalasin D for 2 hrs had a slight but non-significant (n.s.) increase in cilia incidence and increase in mean length (significant at 0.5 µM). Cells treated for 16 hrs had significant increases in cilia incidence and mean cilia length. Minimum of 20 cilia measured per replicate condition. Statistical significance was calculated with a one-way ANOVA with a Dunnett's multiple comparisons test (cilia incidence) and a Kruskal-Wallis test with Dunn's multiple comparisons test (cilia length) as indicated (* $p<0.05$, **=p<0.01, ***=p<0.001, ****=p<0.0001). # indicates the control to which all data-sets were compared.

**(b)** Pharmacological inhibition of non-muscle myosin II ATPase activity through treatment with blebbistatin increased ciliogenesis. hTERT RPE-1 cells stably transfected with LifeAct-GFP were treated for either 2 hrs (upper panels) with blebbistatin or vehicle control after 24 hrs of serum starvation, or for 16 hrs (lower panels) with blebbistatin or vehicle control in serum starvation media (all conditions n=3). Cells treated for 2 hrs had no significant changes in cilia incidence across all blebbistatin concentrations, although there was a significant increase in mean cilia length for concentrations ≥ 2.5 μM. Minimum of 25 cilia measured per replicate condition. Statistical significance was calculated as for (a).

#### Suppl. Figure 7

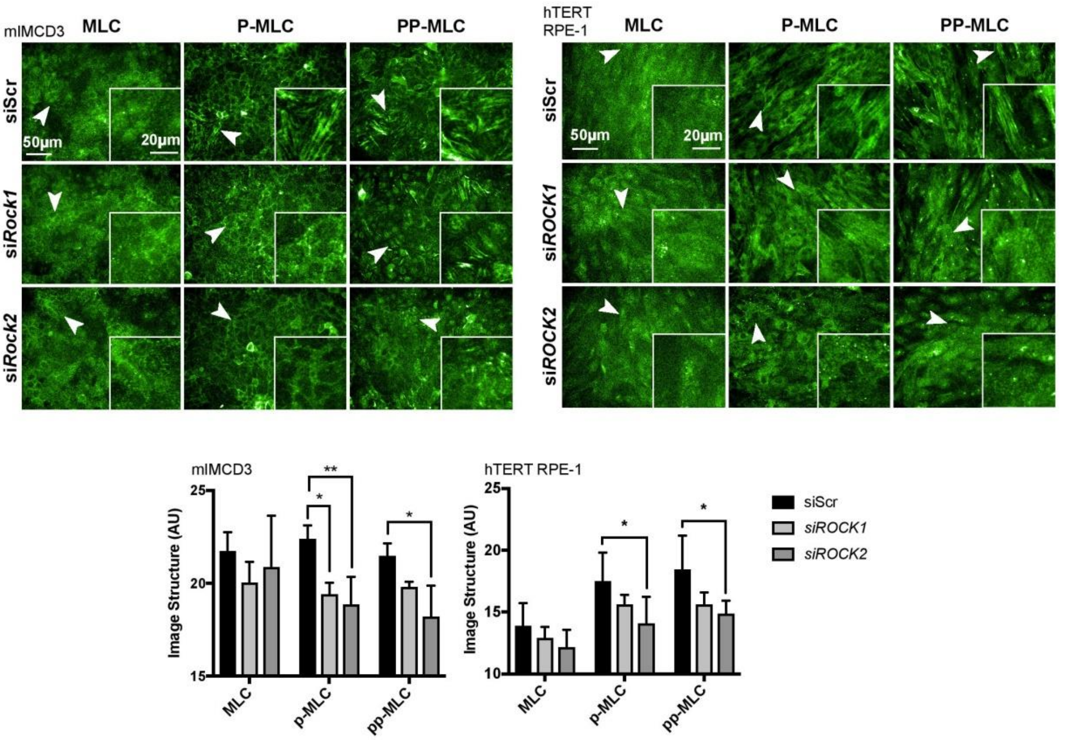


**Supplemental Figure 7: ROCK2 but not ROCK1 knockdown disrupts acto-myosin contraction.**

siRNA knockdown of *ROCK2* inhibits the kinase activity of ROCK2 changes non-muscle myosin IIA organization, visualized by myosin light chain (MLC) staining and the presence of MLC-associated acto-myosin structures in both mIMCD3 (left panels) and hTERT RPE-1 (right panels) cell-lines. Cells were knocked down for with si*Rock1* / si*ROCK1*, si*Rock2* / si*ROCK2* or scrambled siRNA (siScr). Antibodies marked MLC, p-MLC (mono-phosphorylated MLC) and pp-MLC (biphosphorylated MLC at Thr18 & Ser19), with representative immunofluorescence confocal microscopy images shown. p-MLC and pp-MLC visualize acto-myosin fibre-like structures in both mIMCD3 and hTERT RPE-1 cells indicated by arrowheads and displayed in magnified insets. Automated high content image analysis of image texture quantified both p-MLC and pp-MLC fibre-like structures, with bar graphs showing significant decreases for pp-MLC staining following si*Rock2* / si*ROCK2* knockdowns only. Statistical significance was calculated with a two-way ANOVA with Dunnett's multiple comparisons test as indicated (* $p$<0.05, ** $p$<0.01).
